# Detecting differential transcript usage in complex diseases with SPIT

**DOI:** 10.1101/2023.07.10.548289

**Authors:** Beril Erdogdu, Ales Varabyou, Stephanie C. Hicks, Steven L. Salzberg, Mihaela Pertea

## Abstract

Differential transcript usage (DTU) plays a crucial role in determining how gene expression differs among cells, tissues, and different developmental stages, thereby contributing to the complexity and diversity of biological systems. In abnormal cells, it can also lead to deficiencies in protein function, potentially leading to pathogenesis of diseases. Detecting such events for single-gene genetic traits is relatively uncomplicated; however, the heterogeneity of populations with complex diseases presents an intricate challenge due to the presence of diverse causal events and undetermined subtypes. SPIT is the first statistical tool that quantifies the heterogeneity in transcript usage within a population and identifies predominant subgroups along with their distinctive sets of DTU events. We provide comprehensive assessments of SPIT’s methodology in both single-gene and complex traits and report the results of applying SPIT to analyze brain samples from individuals with schizophrenia. Our analysis reveals previously unreported DTU events in six candidate genes.

## Introduction

Alternative splicing enables eukaryotic cells to produce a diverse batch of transcripts and, consequently, proteins from a single gene. While for some genes these distinct transcripts (isoforms) may be used interchangeably, many protein-coding genes have a dominant isoform that is favored in expression across the healthy individuals of a human population.^1^ Predominant expression of alternative isoforms may subject these genes to changes and potential errors in their function.^2^ Differential transcript usage (DTU) analysis is conducted using RNA-Seq data to search for systematic differences in the expression ratios of isoforms that may explain changes in phenotype between cell types, tissues, or populations^2, 3^.

Isoform abundance is often tissue-specific, and DTU (also called isoform switching) may result in proteins with distinct functions, which in turn may play different roles in the cell.^2-6^ There is also a growing interest in the effects of DTU in complex human diseases. Instances of DTU have been associated with DNA repair, numerous human cancer types, heart failure, and psychiatric diseases such as autism, schizophrenia, and bipolar disorder.^7-9^ State-of-the-art DTU analysis tools provide a framework to detect cases where the isoform proportions are consistent within and significantly different between any two groups of samples. However, transcriptomic profiles within populations comprising individuals affected by a complex disease are rarely consistent due to a multiplicity of causal events and disease subgroups; i.e., a cohort of patients diagnosed with the same disease might actually have several distinct underlying genetic disorders.^10^ Therefore, a DTU analysis method that measures and accounts for the structured heterogeneity within complex disease populations is still needed.

We present SPIT, a statistical tool that identifies subgroups within populations at the transcript level and compares their isoform abundance measures. Using both simulated and real RNA-Seq data from human heart tissue, we show that SPIT improves specificity rates compared to the state-of-the-art tools with similar sensitivity, and detects DTU events exclusive to subgroups as well as DTU events shared amongst all case samples. Downstream of DTU analysis, SPIT uses detected DTU events to provide insight into potentially hierarchical subgrouping patterns present in complex disease populations using hierarchical clustering.

Within the SPIT algorithm, subgroups with divergent abundances for each transcript are detected using a kernel density estimator, after which the distributions are compared via a nonparametric Mann-Whitney *U* test. SPIT provides a conservative approximation of the biological and technical variability within datasets with its SPIT-Test module, significantly reducing false-discovery rates. Rather than estimating the expression variability per transcript, SPIT-Test samples a null distribution of minimal *U* statistic *p*-values based on the control group and assumes that, for each transcript, the minimal *U* statistic *p*-value is drawn from the same underlying distribution when there is no real disease association independent of biological or technical variability.

We applied SPIT to search for DTU events associated with schizophrenia, a psychiatric disorder canonically recognized as a heritable complex disease with an undetermined number of subtypes.^11-13^ Genetic causes of schizophrenia have long been studied, however, a clear consensus on the level of genetic liability or the acting set of causal events has not been reached to this day. Whole genome, exome and RNA sequencing studies suggest that a wide range of both common and rare genetic variations, including single-nucleotide polymorphisms (SNPs), copy-number variations (CNVs), ultra-rare coding variants (URVs), and alternative splicing events, may contribute to the pathogenesis of schizophrenia.^9, 14-16^ After analyzing RNA-Seq data from the dorsolateral prefrontal cortex (DLPFC) of 146 schizophrenia patients and 208 controls, SPIT identified six candidate genes that had statistically significant DTU events associated with schizophrenia. Previously-reported disease associations for these candidate genes include neurodegenerative and psychiatric disorders such as Alzheimer’s disease, bipolar disorder, schizophrenia, major depressive disorder, attention-deficit hyperactivity disorder, and autism spectrum disorder. No previous report had identified DTU events in any of these genes.

SPIT is open-source software freely available at *https://github.com/berilerdogdu/SPIT*. Additionally, a user-friendly Google Colaboratory configuration and step-by-step guide are provided at *https://colab.research.google.com/drive/1u3NpleqcAfNz_0EAgO2UHItozd9PsF1w?usp=sharing*.

## Results

### A demonstration on simulated data

A DTU event is defined as a significant difference in the proportions of isoforms contributing to the overall expression of a locus between individual or groups of samples. We are particularly interested in cases where there is a clearly dominant isoform in healthy individuals, where DTU can potentially disrupt cellular function and cause anomalies.

We describe a modeled DTU case with artificially generated data in order to exemplify such DTU events, and to demonstrate the key steps of the SPIT algorithm. Consider a locus from which two distinct isoforms, Isoform 1 and Isoform 2, are transcribed as represented in Figure 1.a. Suppose that the protein translated from Isoform 1 is a functional protein, whereas Isoform 2 gets translated into a dysfunctional, aberrant protein. Consequently, the primary expression profile of this locus in a healthy individual is expected to be Isoform 1. Figure 1.b shows the relative abundances of Isoform 1 and Isoform 2 for four individuals with varying levels of expression at the locus. The left panel of Figure 1.b demonstrates a clear example of DTU between Individual 1 and Individual 2, with Isoform 1 dominant for Individual 1 and Isoform 2 dominant for Individual 2. The right panel of Figure 1.b illustrates why changes in overall expression at the gene/locus or transcript/isoform level are not sufficient indicators of DTU, as illustrated for the same isoforms in Individuals 3 and 4, where overall expression changes but the relative proportion of the isoforms remains the same.

**Figure 1:**
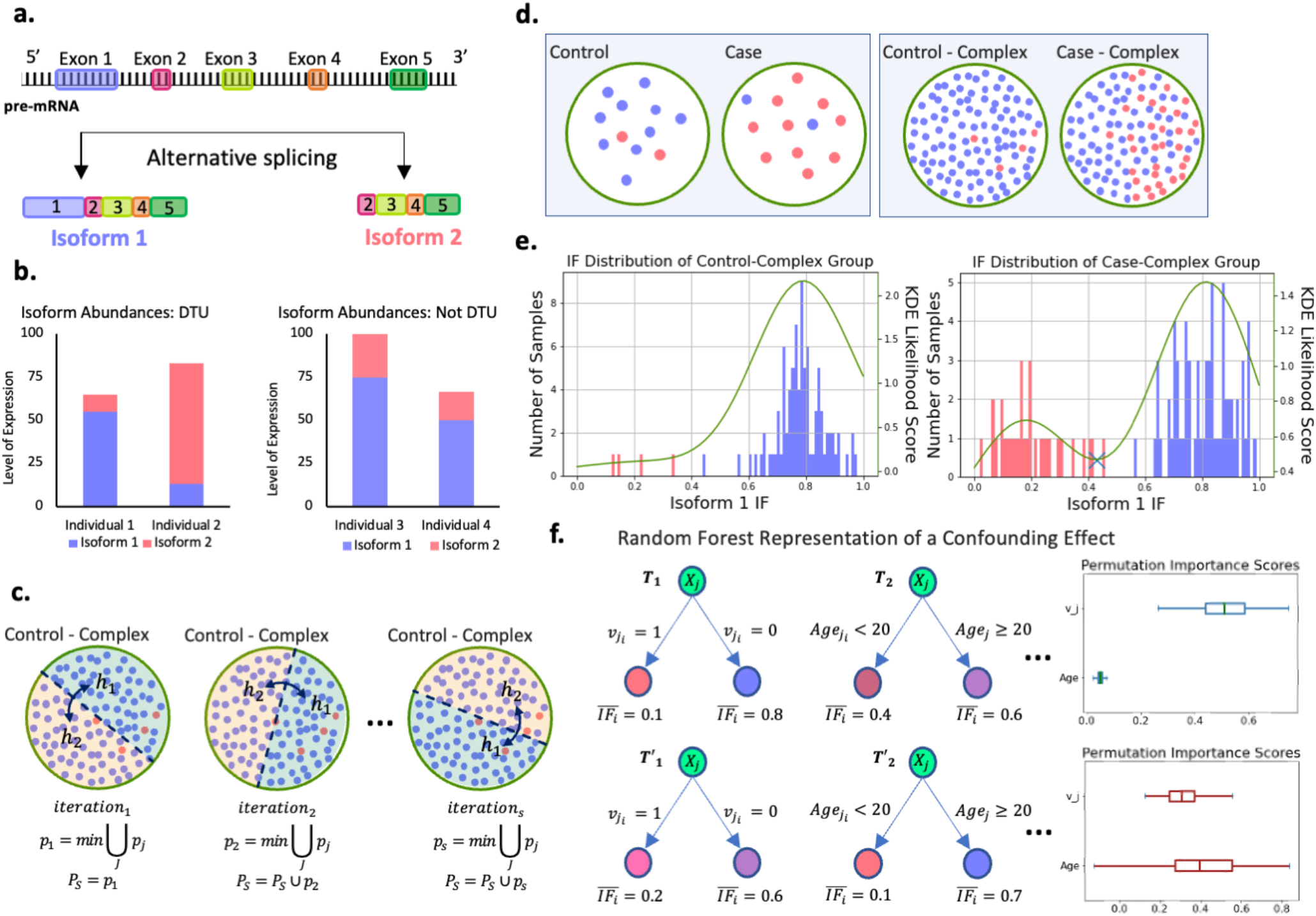
DTU detection demonstration. **a**. Gene locus going though alternative splicing to produce Isoform 1 and Isoform 2. **b**. Left panel: Isoform abundances in a sample case of DTU between individuals 1 and 2. Right panel: Isoform abundances in a sample case without DTU but with changes in overall expression between individuals 3 and 4. **c. Three** SPIT-Test iterations demonstrated with random splits of the Control-Complex group. Samples (dots) are color coded based on their dominant isoforms for the locus in c-f, with blue=isoform 1 and red=isoform 2. **d**. Left panel: Conventional DTU analysis assumption with no structured heterogeneity in either group. Right panel: Heterogeneity structure in complex disease samples, where a subset of cases share the same genetic abnormality (Case-Complex). **e**. Corresponding isoform fraction (IF) distributions for the samples represented in groups Control-Complex and Case-Complex. **f**. Random forest regression representation when there is not a significant confounding effect in the DTU transcript (Upper panel) vs. when there is a clear confounding effect by the covariate “age” (Lower panel). Corresponding permutation importance scores for age and *ν*_*i*_ are shown on the right.

DTU analysis usually entails comparing two groups of samples rather than individuals. In the interest of brevity, suppose for any given individual, either Isoform 1 or Isoform 2 is significantly dominant for the locus in our model DTU case, and note that each individual is color-coded based on their dominant isoform in Figure 1.c-e. Small sample sizes are quite common in RNA-Seq experiments^17^, and the left panel of Figure 1.d represents a typical experiment setup for DTU analysis with 12 samples in each group. If a DTU event between Isoform 1 and Isoform 2 has a causal link to a disease, the left panel of Figure 1.d depicts the expected scenario for a simple genetic disease where the disease is caused by a single or a small set of genes. In this scenario, one assumes that all or nearly all controls have normal gene expression patterns, while the cases all share a distinct but abnormal gene or transcript expression pattern that has caused them to be placed in the disease cohort.

In contrast, the causal set of genes or events are not expected to be shared amongst all individuals affected by a complex disorder. The idea that the majority of complex disorders are likely polygenic, and that distinct combinations of causal events might lead to similar pathogenesis in different patient groups is widely accepted.^18^ When focusing on a particular causal event such as the DTU case between Isoform 1 and Isoform 2, this implies that only a subgroup of patients within the case group are likely to have this event among their causal factors, as depicted in the right panel of Figure 1.d. By segregating this subgroup from the remaining case group, we gain the capability to detect a DTU event that might have otherwise gone unnoticed, and to differentiate potential subclusters of the disease group based on shared DTU events.

In order to do so, we compare the distributions of isoform fractions (*IF*s) between the two groups, which refers to the proportion of total expression attributed to each isoform. Figure 1.e shows the *IF* levels for Isoform 1 in both Control-Complex and Case-Complex groups, which is expectedly high for individuals with Isoform 1 as the dominant isoform at the locus, and low for individuals with Isoform 2 as the dominant isoform. By fitting a kernel density estimator (KDE)^19-21^ on the *IF* distributions, we can search for bimodality, which if found indicates a separation within the groups themselves. The right panel of Figure 1.e demonstrates the clear partition of the Case-Complex subgroups by a global minimum marked with a cross on the KDE curve. We should note that SPIT does not presuppose the existence of a partition in populations and still detects any shared DTU events in the absence of bimodality.

### Partitioning of subgroups

The transcript counts are transformed into *IF*s for each sample as follows:

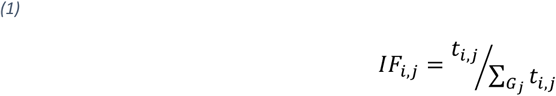

where *IF*_*i,j*_ is the isoform fraction for transcript *j* in sample *i, t*_*i,j*_ is the transcript count for transcript *j* in sample *i*, and *G*_*j*_ stands for the set of all transcripts that belong to the same gene as transcript *j*. We fit a KDE with Gaussian kernel^19-21^ (details on bandwidth selection are described in the Methods section on parameter fitting) on the two vectors of 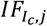, where *I*_*c*_ stands for the samples in groups *c* ∈ {case, control}. If the 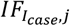 distribution is bimodal, indicating a significant stratification of two subgroups based on the dominance status of transcript *j*, we observe this as a global minimum of the KDE (Figure 1.e). While we acknowledge the possibility of observing a similar divergence within the control group due to technical or biological variability, our primary objective is to identify subgroups within the case samples for potential associations with disease status. The KDE on control group is utilized for flagging the most significant candidate DTU genes as described in the Methods section.

There are several advantages to detecting subgroups based on density estimation, the most important of which is the ability to avoid an underlying distribution assumption for the data set, which can be challenging for RNA-Seq driven data even after multiple normalization steps.^22^ Furthermore, while outlier samples can alter the shape of a KDE, they have a relatively negligible impact on the global minima/maxima as long as appropriate smoothing is applied.^21^ Unlike *k*-means or hierarchical clustering methods, there is not a hyperparameter that fundamentally effects whether or not clusters are detected in the data, and the choice of the bandwidth parameter (*h*) works in our advantage to account for overdispersion by oversmoothing (see Methods section on parameter fitting).

In the presence of a global minimum in the case group at *IF*_*i,j*_ = *m*_*case*_, we define the left tails of the case and control *IF*_*j*_ distributions as the samples that fall to the left of point *m*_*case*_, and the right tails as the samples that fall to the right:

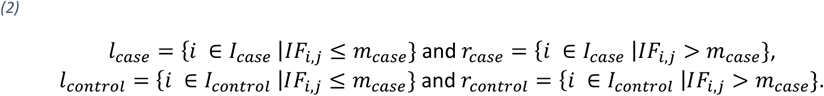

To search for candidate DTU events in *l*_*case*_ and *r*_*case*_ independently, the left tails of the case and control *IF*_*j*_ distributions are compared internally, as are the right tails, using the non-parametric Mann-Whitney *U* test. I.e. 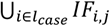 is compared with 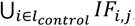, while 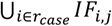 is compared with 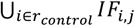. This analysis determines whether the samples in *l*_*case*_ could have been drawn from the left-tail control samples with *IF*_*i,j*_ ≤ *m*_*case*_, or if they exhibit significant differences. Likewise, the same rationale applies for the right tails.

In the absence of a global minimum, a Mann-Whitney *U* test is conducted between the entire groups of *I*_*case*_ and *I*_*control*_.

### Estimating dispersion with SPIT-Test and detecting DTU

Although the use of non-parametric statistical tests can help control the false discovery rate (FDR) in differential analyses, the effectiveness of several competing methods is notably diminished when the input data is overdispersed with outliers^23^, a common characteristic of RNA-Seq data^24^. This prevalent phenomenon suggests that we are not capable of precisely estimating dispersion for each individual transcript or gene, in addition to not being able to adequately correct for the vast number of hypotheses being tested. To overcome this challenge, we choose to estimate a single null distribution for the minimal Mann-Whitney *U*-statistic *p*-values, and assume that these observed minimal *p*-values reflect the upper threshold of dispersion in the input dataset.

The true null distribution *P*_*S*_ of the minimal *U*-statistic *p*-values represents the lowest expected *p*-values when there is no real association between the phenotype of interest, such as a disease, and the changes in isoform dominance among individuals or groups. To estimate 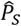, SPIT-Test takes advantage of the control group in which disease association is absent, yet individual differences due to biological, technical, or other confounding factors can be observed. As illustrated in Figure 1.c, SPIT-Test is an iterative process which randomly splits the control group in half, and identifies the most significant difference in isoform fractions between the two halves. Later on, the candidate DTU events between the case and control groups are compared, in terms of their significance, to the observed differences between random halves of the control group.

The following steps are performed at each iteration *s*:

1. Randomly split the control samples into two sets of equal size, namely *h*_*k,s*_ where *k* ∈ {1, 2} represents each half for iteration *s*.
2. Select a random split point *o*_*s*_, to define the left and right tails of each half as:

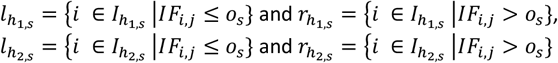
3. For each transcript *j*, conduct a Mann-Whitney *U* test between the sets of 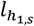 and 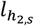, yielding a Mann-Whitney *U*-statistic *p*-value 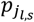. Similarly, conduct a Mann-Whitney *U* test between the sets of 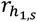 and 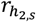, yielding 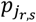.
4. Assign 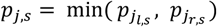 to each transcript *j* for iteration *s*.
5. Among the *U*-statistic *p*-values assigned to all transcripts, store *p*′_*s*_ = min ⋃_*J*_ *p*_*j,s*_. In order to avoid excessive influence from outlier transcripts, we only sample *p*′ once from the same transcript throughout all iterations. In other words, in iteration *s* we consider transcripts from which 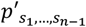 have not been sampled.
6. 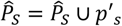.

SPIT-Test estimates dispersion on a global scale, assuming that any transcript could have been subject to the highest observed level of dispersion. Therefore, for an arbitrary transcript *j*, 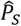 is considered as an empirical null distribution of the minimal *U*-statistic *p*-value. This approach emulates the min-P and max-T procedures^25^, and is employed to set a *p*-value threshold, *p*′_*threshold*_, based on 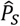 that determines the set of candidate DTU transcripts between case and control samples as:

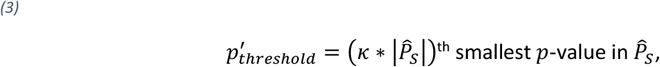

where *κ* is a user-set parameter. For instance, if *κ* = 0.1 for 1000 iterations, the threshold would be the 100th smallest *p*-value. SPIT-Test deviates from a traditional permutation test in its randomization steps 1 and 2, and its exclusion of the case samples due to the potential presence of unknown subgroups. Although *κ* cannot directly translate into a target family-wise error rate (FWER), we experimentally show that smaller values of *κ* achieve remarkable control over FWER.

### DTU simulation and evaluation

Simulated RNA-Seq reads are conventionally used to evaluate differential analysis tools, as we lack knowledge of ground truth in real data. However, research has consistently shown that simulated reads do not accurately represent the overdispersion levels in real RNA-Seq experiments, leading to underestimation of FDR.^23, 26^ In order to obtain a more accurate assessment of SPIT’s performance, we make use of both simulated and real RNA-Seq data. In these two types of evaluation sets, we compare the true positive rate (TPR) and FDR outcomes of SPIT, and the state-of-the-art tools *DEXSeq*^27^ and *DRIMSeq*^28^ used together with the stage-wise adjustment tool *stageR*.^29^

### Evaluation with simulated RNA-Seq reads

We borrow the DTU simulation with the largest sample sizes from the “Swimming Downstream” pipeline by Love *et al*.^30^ as our test dataset with simulated RNA-Seq reads. (Please see the corresponding Methods section for details.) This dataset simulates a large number of (> 1500) DTU events in relatively homogenous populations, resembling the scenario depicted in the left panel of Figure 1.d. While dispersion is incorporated into the transcript expression patterns, there are no subgroups or divergence in the DTU events.

The TPR and FDR at the gene level are reported for each tool in Figure 2.b, where both *DEXSeq* and *DRIMSeq* have 3 outcomes corresponding to *stageR* target overall FDR (OFDR) values 0.01, 0.05, 0.1. For SPIT, we report 5 outcomes corresponding to setting hyperparameter *κ* = 0.2, 0.4, 0,6, 0.8 and 1 on 1000 iterations. Although the tuning of target OFDR for *stageR* and *κ* for SPIT are not directly comparable, lower values of both parameters lead to more conservative behavior, allowing better control over FDR and often yielding decreased TPR.

**Figure 2:**
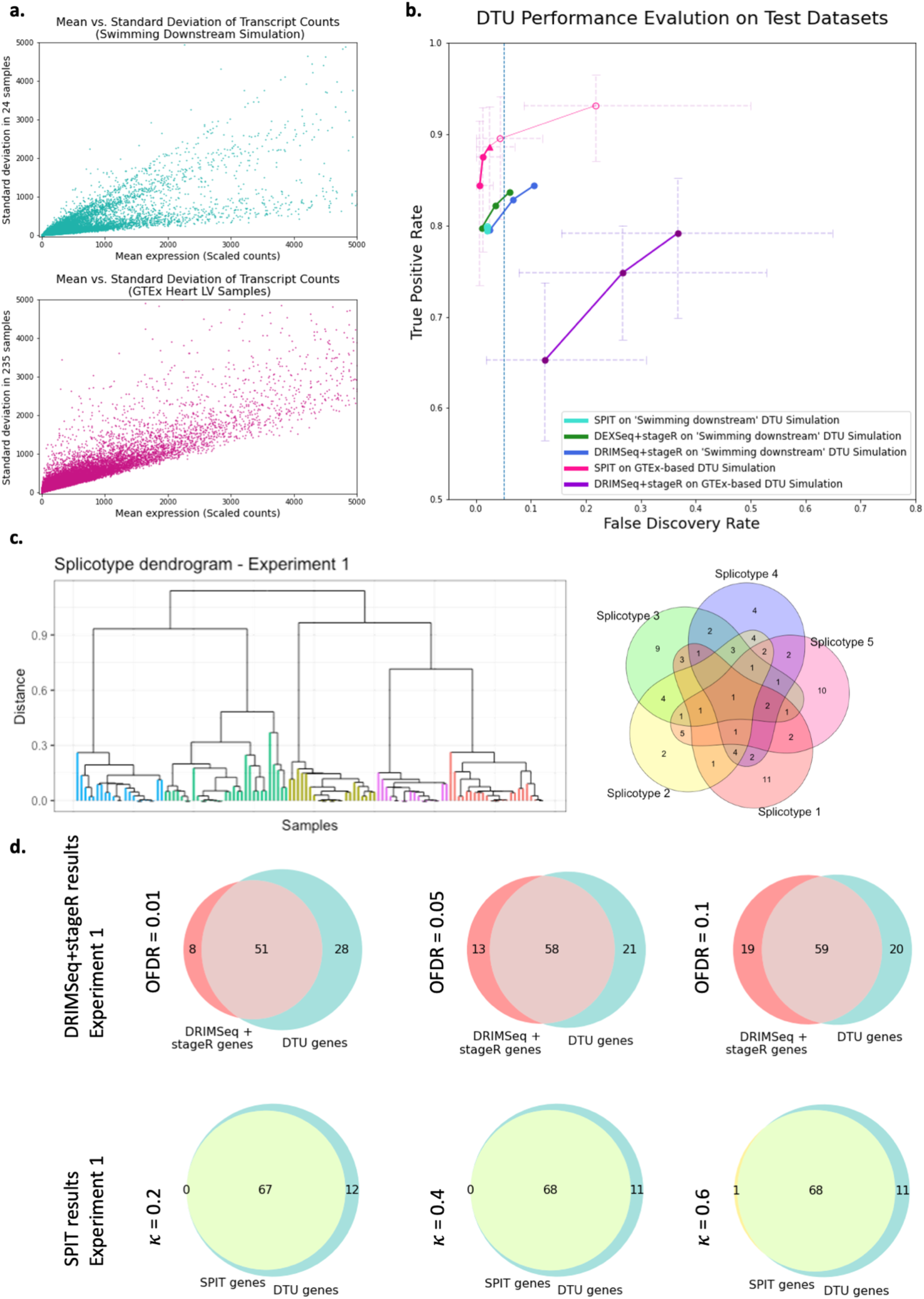
**a**. Mean vs. standard deviation of the transcript counts are plotted for the Swimming Downstream and GTEx experiment samples to represent relative dispersion levels. **b**. Gene-level DTU-performance evaluation on both Swimming Downstream and GTEx test datasets. Radical values of *κ* = 0.8 and *κ* = 1 are included (as unfilled circles) to show the effects of hyperparameter adjustment. **c**. The DTU event sharing Venn diagram for the first experiment in the GTEx simulations (Right), and the corresponding final subcluster dendrogram based on the SPIT DTU matrix (Left). The subclusters are color coded based on their distinct sets of simulated DTU events (splicotypes). **d**. Overlap of DRIMSeq+stageR pipeline (Top) and SPIT (Bottom) results with simulated DTU genes in the first experiment.

TPR and FDR outcomes of *DEXSeq* and *DRIMSeq* were consistent with the “Swimming Downstream” evaluation. Both tools yielded high sensitivity levels while *DEXSeq* maintained a slightly better control over FDR. On the same simulated dataset, SPIT yielded a comparable yet slightly lower TPR value while always keeping the FDR lower than 0.05. Different values of the hyperparameter *κ* did not result in noticeable differences in TPR or FDR on this dataset.

### Evaluation with real RNA-Seq reads

To form the basis of our test dataset with real RNA-Seq reads, we quantified Illumina reads of 235 normal heart (left-ventricle) samples obtained from the Genotype-Tissue Expression (GTEx) project^31^. Figure 2.a shows the mean-standard deviation plots of the two datasets, revealing a significantly higher level of dispersion in the GTEx dataset compared to the “Swimming Downstream” dataset of simulated RNA-Seq reads.

Next, we conducted 20 separate experiments in each of which we compared random halves of the GTEx dataset after introducing 100 simulated DTU events into one of the halves (please see the corresponding Methods section for details). In an effort to model the expected heterogeneity in a complex disease group, we distributed the 100 DTU events between 5 subgroups in such a way that some DTU events are shared between the subgroups while some are exclusive (see Figure 2.c for an example). For the rest of the paper we’ll refer to any such subgroup that shares the same DTU events as a “splicotype” group.

In any random partition of real RNA-Seq samples into two groups, it is not certain that there are no actual DTU events beyond the ones we introduced. Therefore, the TPR and FDR measures for the GTEx experiments are only estimates. Our hypothesis in evaluating these experiments was that if any method consistently detected additional DTU events between random partitions of a healthy sample group, the discoveries were either noise or else due to biological variance that are not of interest. As such, we present the mean estimated FDR and TPR values of 20 experiments for SPIT and *DRIMSeq*+*stageR* pipeline in Figure 2.b with error bars indicating the minimum and maximum FDR/TPR values obtained. Additionally, in Figure 1.d, we show the individual Venn diagrams representing the overlap between the *DRIMSeq*+*stageR* pipeline and SPIT results with the simulated DTU genes for the first experiment out of the 20 conducted. Venn diagrams for the remaining experiments showing similar results are provided in Supplementary Fig.1.

Due to its generalized linear model (GLM) fitting step, *DEXSeq* requires significant compute time for large sample sizes. After running for 168 hours on 24 cores and 256 GB RAM, dispersion estimation for the first experiment remained unfinished. Therefore, we only compare *DRIMSeq* + *stageR* and SPIT results for the GTEx experiments.

In line with the “Swimming Downstream” evaluation, we applied *DRIMSeq* followed by *stageR* with target OFDR values of 0.01, 0.05, and 0.1 to the GTEx experiments. Because the SPIT pre-filtering process is included in the DTU simulation, we performed *DRIMSeq* + *stageR* analysis on the SPIT-filtered counts and bypassed *DRIMSeq* filters.

In contrast to the TPR and FDR values obtained with the simulated “Swimming Downstream” dataset, the *DRIMSeq* + *stageR* pipeline yielded a wider range of estimated TPR and FDR values on the GTEx experiments. For the GTEx experiments, the *DRIMSeq* + *stageR* pipeline produced lower TPR and notably higher FDR estimates for all target OFDR values (0.01, 0.05, and 0.1), with a more significant difference in performance between each OFDR value. We also note the wide error bars in the pipeline, indicating a large range of performance across all 20 experiments. This variability could be attributed to the distinct biological differences between the random partitions in each experiment or to the level of heterogeneity introduced in the simulation through varying compositions of shared DTU events between random splicotypes.

As with the “Swimming Downstream” dataset, we report TPR and FDR estimates for SPIT obtained by setting hyperparameter *κ* = 0.2, 0.4, 0,6, 0.8 and 1. For input datasets with large number of control samples (*n* ≥ 32), SPIT offers an optional cross-validation procedure to estimate the optimal value *κ*^*^ based on inferred dispersion, which is detailed in the Methods section on parameter fitting. In Figure 2.b, the TPR and FDR obtained using the estimated *κ*^*^ is represented by a triangle, which for this dataset is 0.6. Overall, the estimated TPR and FDR levels for SPIT remained comparable to the values obtained for the “Swimming Downstream” dataset with a slight increase in both TPR and FDR. The gain in sensitivity is expected for SPIT with large sample sizes since it uses the Mann-Whiney *U* test when comparing any two sets of *IF* values. While the optimal *κ*^*^ parameter still has an estimated FDR < 0.05, SPIT’s control over FDR also diminished with real RNA-Seq reads compared to the simulated test set. A clear increase in both TPR and FDR was observed for *κ* = 0.8 and *κ* = 1, which are included to demonstrate the effects of using radically large values for hyperparameter *κ*. The range represented by the error bars in Figure 2.b is smaller for SPIT compared to that of *DRIMSeq*, which indicates higher consistency across all 20 experiments.

Upon detecting the DTU events for any given dataset, SPIT outputs a binary matrix *M* of DTU events that marks the presence (1) or absence (0) of a DTU event at the gene level for any sample in the case group relative to the control group. We show that using SPIT’s output matrix *M*, we are able to cluster the case samples into their separate splicotype groups based on their shared events by applying hierarchical clustering. The chosen distance metric calculates the proportion of unique events between any two samples relative to the total number of DTU events. As shown in Figure 2.c, SPIT perfectly captures the five clusters that were artificially created. Clustering on the first experiment is shown in Figure 2.c based on the SPIT output with *κ*^*^; the remaining experiments are shown in Supplementary Fig.1.

### Detecting known tissue-dependent DTU events

As a positive control experiment, we next investigated a set of four tissue-dependent DTU events that had been previously confirmed individually by various studies and also collectively validated by Reyes & Huber in 2018^32^. For this analysis, we utilized samples from the GTEx dataset (Supplementary Table 1) that were aligned as part of the CHESS 3 project.^33^ Figure 3 visually illustrates differentially expressed transcripts between tissues at each locus. All transcriptional landscape were created using the sashimi plot module in TieBrush after aggregating read alignments from all samples in each tissue. SPIT results on all four DTU events are detailed below.

**Figure 3:**
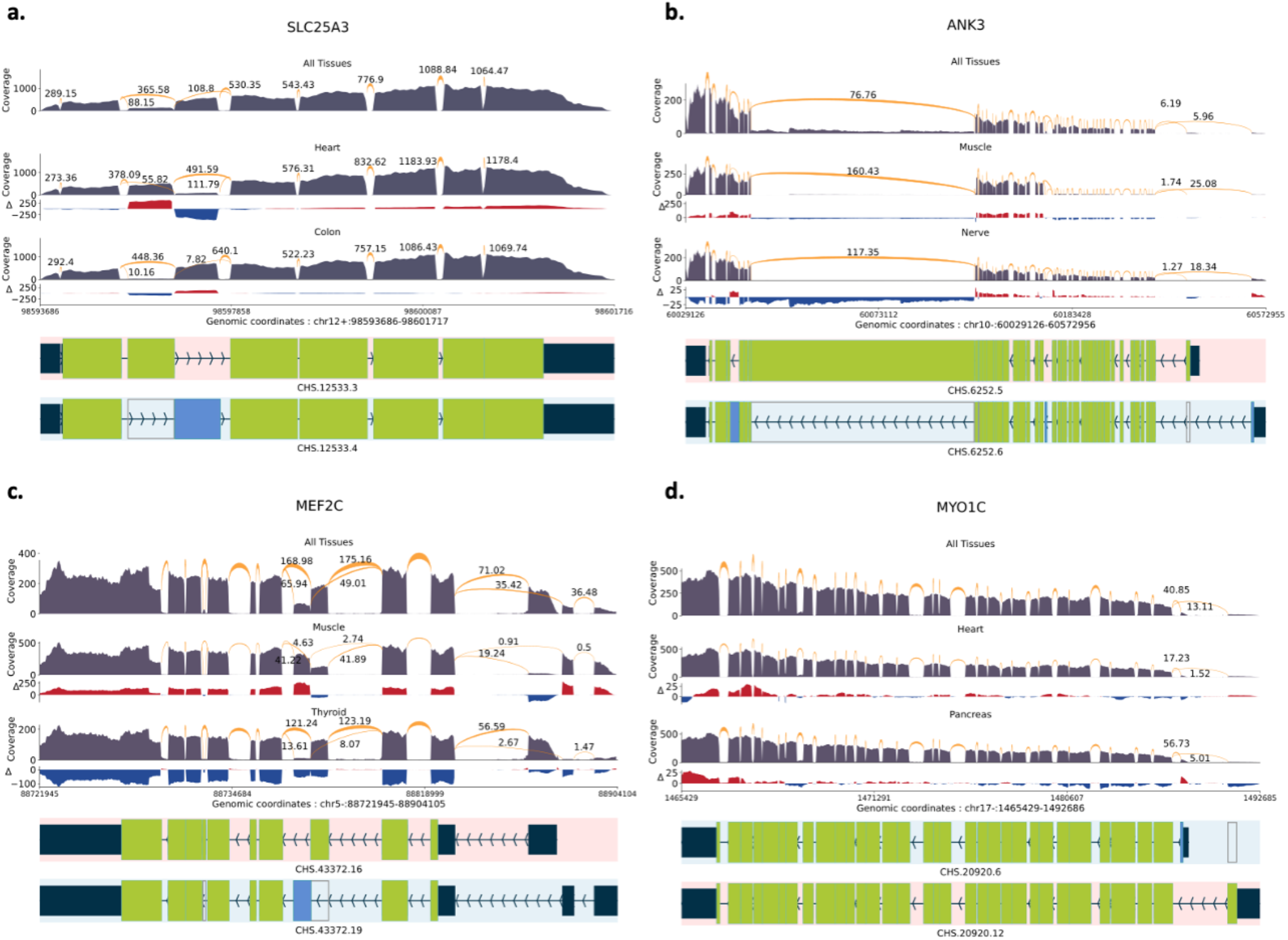
Sashimi plots with normalized coverage and junction values from GTEx samples of CHESS 3 project. Only the relevant isoforms and junction values are included for readability. The normalized coverage values for each tissue were subtracted from the normalized coverage of the entire GTEx dataset, and the results were illustrated as the Δ track. **a**. *SLC25A3* DTU event between heart and colon tissues. **b**. *ANK3* DTU event between muscle and nerve tissues. **c**. *MEF2C* DTU event between muscle and thyroid tissues. **d**. *MTO1C* DTU event between heart and pancreas tissues.

#### SLC25A3

The mitochondrial phosphate transporter gene *SLC25A3* exhibits a phenomenon known as “mutually exclusive exons”^3^, which refers to the observation that specific exons within the gene are spliced into distinct isoforms but they are not simultaneously present within the same isoform. We compared 497 samples of heart tissue and 380 samples of colon tissue from the GTEx dataset, and SPIT was able to confirm that one of these isoforms, which is recognized as the primary expression preference in heart and skeletal muscle, is indeed more prevalent in heart tissue samples (Figure 3.a).

#### ANK3

Together with two more ankyrin genes, *ANK3* plays a crucial role in generating a diverse array of ankyrin proteins in mammals. Tissue-specific splicing of *ANK3* has previously been shown in skeletal muscle and tibial nerve tissues^32, 34^. A total number of 480 muscle and 339 nerve tissue samples from GTEx were analyzed using SPIT, confirming the presence of an isoform switch characterized by alternative start sites and distinct patterns of exon splicing (Figure 3.b).

#### MEF2C

*MEF2* transcription factors are significant in regulating cell differentiation and expression, and they undergo tissue-specific alternative splicing, adding to their functional diversity. *MEF2C* in humans has two mutually exclusive exons, one of which is shown to be more prevalent in skeletal muscle^35^. We compared 480 muscle tissue samples from GTEx with 361 thyroid samples using SPIT and were able to detect the isoform switching as a significant DTU event (Figure 3.c).

#### MYO1C

*Myosin IC* encodes a protein of the myosin family, which serves multiple cellular functions including vesicle transportation, transcription and DNA repair^36, 37^. The presence of a tissue-dependent transcription start site in *Myosin IC* has been demonstrated, leading to splicing of an alternative first exon^36^, which SPIT successfully detects upon comparing 497 heart and 199 pancreas samples from GTEx (Figure 3.d).

### Schizophrenia application

After evaluating its performance, we explore the application of SPIT in identifying DTU genes associated with schizophrenia, where we expect a divergence in the causal mechanisms underlying pathogenesis for individual or groups of patients. We obtained RNA-Seq samples of post-mortem DLPFC tissue from a total of 354 adult brains, which were sequenced by the Lieber Institute for Brain Development.^38^ After applying various quality filtering criteria that are described in detail in the Methods section, we selected 146 schizophrenia samples and 208 control samples for comparison in our analysis (Supplementary Table 2).

The parameter-fitting process was applied to the control samples, resulting in *κ*^*^ = 0.4. Prior to confounding analysis, SPIT detected 135 potential DTU events between the case and control samples. The binary DTU matrix for these 135 transcripts was then inputted to the confounding control module of SPIT which is described in the Methods section. Covariates considered for all samples included sex, race, age, batch identification, and RNA integrity number (RIN) which highly correlates with RNA degradation.^39^ 129 candidate transcripts were eliminated based on their permutation importance scores, leaving a final set of six DTU transcripts in six genes (Figure 4.c). The SPIT-Chart for this analysis (Figure 4.a) shows the relationship between the median *p*-values obtained from 1000 iterations of SPIT-Test and the *p*-values resulting from comparing control and schizophrenia samples for transcripts.

**Figure 4:**
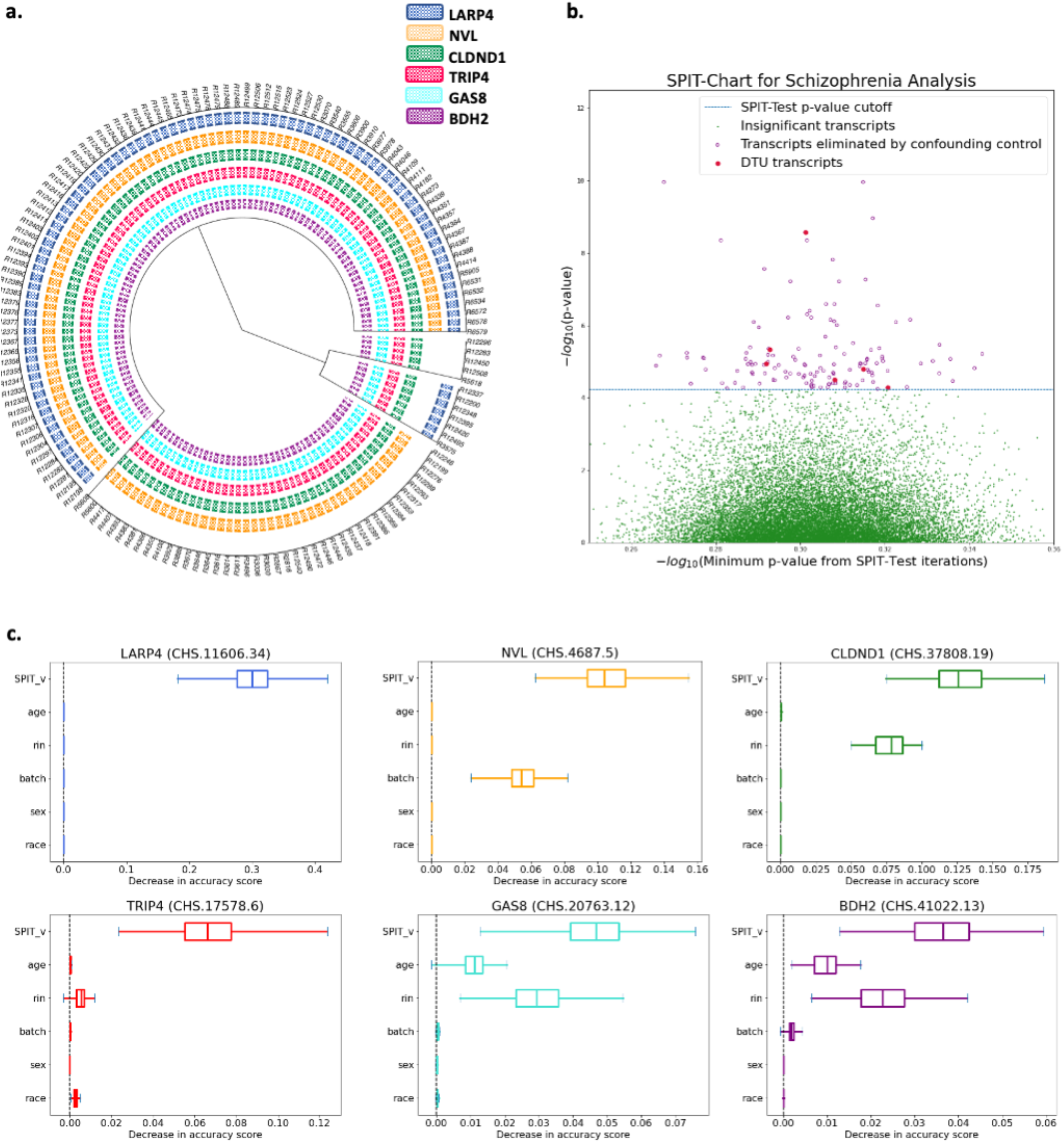
**a**. Dendrogram representation of hierarchical clustering applied on the SPIT DTU matrix for schizophrenia samples **b. SPIT-Chart for the schizophrenia analysis**: For each transcript that passed the initial filtering steps, the median *p*-value that has been observed through 1000 iterations of the SPIT-Test (median (⋃_*S*_ *p*_*i,j*_)) is plotted on the *x*-axis, and the *p*-value observed in the actual comparison of the schizophrenia samples to the controls is plotted on the *y*-axis, both on −*log*_10_ scale. **c**. Box plots of permutation importance scores (generated from 100 permutations) of the SPIT output vector and provided covariates for the final 6 DTU genes.

Amongst the six candidate genes, four (*BDH2, CLDND1, GAS8, TRIP4*) displayed DTU events in all schizophrenia samples, while the other two genes (*LARP4, NVL*) showed significant DTU events in specific subgroups. Figure 4.b depicts the clustering of schizophrenia samples based on identified DTU events, revealing a partitioning into four subgroups in this dataset. We present short descriptions of the functions and associations of the six candidate genes below.

#### GAS8 (Growth Arrest Specific 8)

A multi-tissue study examined SNPs for enrichment of expression quantitative trait loci (eQTL) across 11 genome-wide association studies (GWAS) focused on schizophrenia and affective disorders (including bipolar disorder, major depressive disorder, autism spectrum disorder, and attention-deficit hyperactivity disorder)^40^. The study identified *GAS8* amongst genes affected by the high-confidence cis-eQTLs in multiple brain regions, and reported its cross-disorder associations as well as specific associations with bipolar disorder.

#### NVL (Nuclear VCP Like)

This gene is a member of the AAA family (ATPases associated with diverse cellular activities) and encodes for two proteins with recognized distinct functions, *NVL1* and *NVL2*^41^, the latter of which is involved in regulating ribosome biogenesis in eukaryotes^42^.

There is a growing body of evidence suggesting correlations between disrupted ribosome synthesis and aging, as well as neurodegenerative diseases like Alzheimer’s disease and Parkinson’s disease^43-48^. In the subset of schizophrenia samples where *NVL* is implicated in DTU, we observed that the *NVL1* isoform was preferred, potentially indicating perturbed ribosomal synthesis (Supplementary Figure 3).

#### LARP4 (La Ribonucleoprotein 4)

The protein encoded by this gene enables RNA-binding activity and plays a critical role in translation regulation^49^. *LARP4* has been found to show differential expression between the unaffected siblings and first-degree relatives of schizophrenia patients compared to unaffected individuals unrelated to the patients^50^.

#### BDH2 (3-Hydroxybutyrate Dehydrogenase 2)

This gene is responsible for encoding a siderophore that plays a crucial role in maintaining iron balance within cells, offering protection against oxidative stress^51^. Studies have indicated a significant downregulation of *BDH2* in response to inflammation and endoplasmic reticulum (ER) stress^52^. Disrupted iron homeostasis and ER stress have long been associated with neurodegenerative diseases like Alzheimer’s disease and Huntington’s disease^53, 54^. Recent studies report *BDH2* to be directly implicated in Alzheimer’s disease progression^55^.

#### TRIP4 (Thyroid Hormone Receptor Interactor 4)

The protein encoded by this gene is one of the four components of the activating signal cointegrator 1 (ASC-1) complex. Mutations in ASC-1 components have been described as shared anomalies between the neurodegenerative diseases Amyotrophic Lateral Sclerosis (ALS) and Spinal Muscular Atrophy (SMA)^56^. Mutations in *TRIP4* and *ASCC1*, another component of ASC-1 complex, are widely recognized as a cause of SMA^57, 58^.

#### CLDND1 (Claudin Domain Containing 1)

This gene encodes transmembrane proteins of tight junctions, which play a role in regulating the permeability of brain endothelial cells^59^. *CLDND1* has been linked to Alzheimer’s disease^60^, with one study indicating a potential correlation specifically with a subgroup of the condition^61^.

## Discussion

Transcriptomic profiles in populations with complex diseases can exhibit inherent complexity where differentially expressed events are not necessarily shared among all individuals affected by the specific disorder. Consequently, applying the same statistical assumptions for these populations as those used for simple genetic disorders can lead to misleading results in differential analyses. SPIT is the first DTU tool built to accommodate and detect structured heterogeneity within populations. Through DTU simulations built on GTEx samples, we show that SPIT not only achieves improved sensitivity and specificity in detecting DTU genes in heterogeneous populations, but also successfully captures the specific DTU events for the prevalent subpopulations present.

Our results on the “Swimming Downstream” dataset by Love *et al*. also demonstrate that SPIT is equally effective on relatively homogeneous populations, and proves to be applicable for diverse scenarios, including simple genetic disorders, tissue-to-tissue comparisons, and other types of DTU studies. SPIT consistently maintains notably low false discovery rates regardless of the level of dispersion in the datasets.

In addition to simulated experiments, we present four previously confirmed tissue-specific DTU cases that SPIT successfully detected in GTEx samples, as well as six novel DTU associations with schizophrenia. However, to establish any causal link between these six candidate DTU events and schizophrenia, a much more comprehensive investigation is needed, which is beyond the scope of this paper.

## Methods

### Pre-filtering

The main input SPIT requires is transcript-level count data from an RNA-Seq quantification tool, a mapping file that assigns gene names to each of the transcripts, and any metadata for the samples. Pre-filtering the transcripts before DTU analysis has been shown to improve performance for state-of-the-art tools^30, 62^, which also holds true for SPIT. The default behavior of SPIT involves the stringent pre-filtering steps listed below which build upon the *DRIMSeq* filtering criteria:

1. Each transcript must have a Counts per million (CPM) value of at least 10 in at least *n*_*small*_ samples, where *n*_*small*_ is a user-set parameter that defines the smallest sample size presumed for the subgroups within populations.
2. Each transcript must have a positive read count in at least a fraction *p*_*r*_ of the samples in both the case and control groups, respectively. *p*_*r*_ is a user-set parameter and defaults to 0.20.
3. Each gene must have a read count of at least *g*_*c*_ in at least *g*_*n*_ samples, where *g*_*c*_ and *g*_*n*_ are user-set parameters and default to 10.
4. Each transcript must have an *IF* value larger than *f* in at least *n*_*small*_ samples, where *f* is a user-set parameter and defaults to 0.1.
5. After the filtering steps above, there must remain at least 2 transcripts for each gene.
6. The control group must have a consistently dominant isoform for each gene. This criterion is met for a gene when the same isoform of the gene has the largest *IF* in at least a fraction *p*_*d*_ of the control samples, where *p*_*d*_ is a user-set parameter and defaults to 0.75.

As is the case for any filtering criteria prior to differential analyses, these steps may inadvertently exclude genuine DTU cases and lower sensitivity. Thus, while these steps are included and recommended in the SPIT pipeline, any or all of them can be excluded from the analysis by the user. Supplementary Figure 2 outlines the application of this filtering pipeline on the schizophrenia samples discussed in the Results section.

### Test set with simulated RNA-Seq reads: “Swimming Downstream”

Love *et al*. simulated DTU events in 1,500 genes by swapping Transcript Per Million (TPM) abundance values between two isoforms. In an additional 1,500 genes, they simulated differential transcript expression (DTE) by altering the abundance value of a single isoform by a fold change between 2 and 6. For these DTE genes, if the differentially expressed transcript is not the only isoform, they were also considered DTU cases as the relative isoform abundances were also expected to change. We include both types of these DTU events in our analysis.

Love *et al*. conducted four experiments with various sample sizes in the case and control groups (*n* = 3 vs. 3, *n* = 6 vs. 6, *n* = 9 vs. 9, *n* = 12 vs.12) to evaluate state-of-the-art DTU tools *DEXSeq, DRIMSeq, RATs*, and *SUPPA2*. They reported that while *SUPPA2* and *RATs* always controlled their FDR, their sensitivity levels remained consistently low across all experiments, hovering around 50%. *DRIMSeq* and *DEXSeq* had considerably higher sensitivity (≥ 75%) while sometimes exceeding their target FDR. Both *DRIMSeq* and *DEXSeq* demonstrated improved FDR control with larger sample sizes, and 12 vs. 12 yielded the most favorable TPRs and FDRs.

Based on these findings, we choose to reproduce the “Swimming Downstream” results obtained with *DEXSeq* and *DRIMSeq* on the *n* = 12 vs. 12 experiment and to evaluate SPIT’s performance on the same dataset. We first ran *DEXSeq* and *DRIMSeq* on the released Salmon ^63^ quantification files by Love *et al*.^64^ as outlined in the “Swimming Downstream” workflow. We then applied the stage-wise adjustment tool *stageR* on the preliminary results from both *DEXSeq* and *DRIMSeq* using target OFDR values 0.01, 0.05, and 0.1. The “Swimming Downstream” evaluation first applied the *DRIMSeq* pre-filters on the simulated counts and defined the set of true positives as the DTU genes and transcripts that pass the *DRIMSeq* filter. To be able to replicate the reported TPR and FDRs and compare results, we applied the same filters and skipped the SPIT pre-filtering process.

### Test set with real RNA-Seq reads: GTEx simulation

To simulate each of the 20 GTEx experiments the following steps were executed:

1. Randomly divide the 235 GTEx samples into two sets to create case and control groups, *I*_*case*_ and *I*_*control*_, comprising of 117 and 118 samples, respectively.
2. Apply the SPIT pre-filter outlined above assuming the randomly assigned *I*_*case*_ and *I*_*control*_. Note that we skip step 6 of the pre-filtering as it could create an unfair bias in the pre-filtered set of genes towards the DTU genes selected in the next step.
3. We apply the criteria outlined in step 6 of the pre-filtering process to identify genes with consistently dominant isoforms within the *I*_*control*_ group. Out of these genes with dominant isoforms, we randomly select 100 to compose our superset of DTU genes, D = {d_1_, d_2_, …, d_100_}.
4. For each splicotype group (subgroup of samples that share the same DTU events) *π*_*s*_, *s* ∈ {1, 2, 3, 4, 5}, we randomly select 30 DTU genes from set *D* with replacement to form 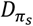. This results in a complex and structured partition within *I*_*case*_, where some DTU genes are shared between the five splicotypes while others are unique to a specific splicotype.
5. For a DTU gene 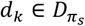, let *α*_*k*_ be the dominant isoform of *d*_*k*_ in *I*_*control*_ with 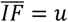, and *β*_*k*_ be the least dominant isoform in *I*_*control*_ with mean 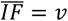. We switch the dominance status of *a*_*k*_ and *β*_*k*_ in *I*_*case*_ by allowing 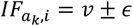 and 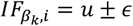 for all 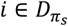, where noise parameter *ϵ* = 0.05.
6. Within all simulated DTU cases, the original transcript counts for *a*_*k*_ and *β*_*k*_ are updated by multiplying the gene counts by 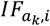 and 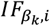, respectively. The gene counts are updated subsequently as the sum of all updated transcript counts, and *IF* values are calculated once again with equation (1) so that within each gene *IF* values add up to 1.

### Addressing confounding variables

After completing the preliminary DTU analysis, the main output of the SPIT pipeline is a binary vector *ν*_*j*_ for each transcript indicating the presence (1) or absence (0) of a DTU event in each sample in comparison to the control group. Note that *ν*_*j*_ carries a 0 for all samples of the control group. Moreover, notice that for the transcripts that SPIT reports as significant DTU events, the *ν*_*j*_ vector represents a partitioning of all samples, case and control, into two groups with relatively high and low *IF*_*j*_ values.

In the presence of a confounding effect, this partition of the high and low *IF*_*j*_ values can also be achieved via the confounding variable if included in the experimental design. Based on this assumption, SPIT filters out the DTU events with potential confounding effects using a random-forest-based method.

Given a set of covariates *X* = {*x*_1_, *x*_2_, …, *x*_*k*_}, we define a matrix *X*_*j*_ for every candidate DTU transcript *j* such that 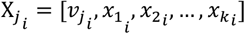 for any sample *i* in either group. We also define a vector *y*_*j*_ based on the *IF*_*j*_ values such that 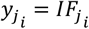 in the same sample order as in *X*_*j*_.

We then fit a random forest regressor^65, 66^ *ϕ*_*j*_(*X*_*j*_) → *y*_*j*_ on each candidate DTU transcript. The same number of samples as in the input matrix is bootstrapped for the construction of each tree with maximum tree depth 1, and we minimize the *l*_2_ loss on the mean *IF*_*j*_ in terminal nodes. Notice that with tree-depth 1, our goal is not to precisely predict 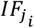 for samples as much as it is to assess which covariates might be contributing into observable variance in *IF*_*j*_ values. We require at least *n*_*small*_ number of samples to split the root node. An illustrative case of detecting a confounding effect can be seen in the random forest depicted in Figure 1.d. Building on the modeled demonstration in Figure 1, assume that a candidate DTU event was detected for the subgroup in Case-Complex samples. Supposing one covariate (age) was provided as input, the random forest attempts to regress *IF*_*j*_ based on 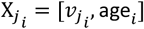. On the upper panel, the first tree *T*_1_ finds the expected effectiveness of vector *ν*_*j*_ in separating low *IF*_*j*_ values, as it was primarily inferred based on *IF*_*j*_. A similar effective partition cannot be achieved with the provided covariate age in tree *T*_2_.

On the lower panel, however, a partition by age in *T*′_2_ demonstrates that age works as well as *ν*_*j*_ in *T*_1_, which implies that the identified DTU event cannot be confidently distinguished from a possible confounding effect of the covariate.

With the objective of estimating the importance of each covariate as well as *ν*_*j*_ in the partitioning of high vs. low *IF*_*j*_ samples, we conduct a permutation importance test^66, 67^ on each random forest *ϕ*_*j*_. The permutation importance test is based on the coefficient of determination 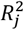 of *ϕ*_*j*_, which is a score of how well *IF*_*j*_ is predicted in tree leaf nodes.

Let *ϕ*_*j*_ have *l* leaf nodes *λ*_1_, …, *λ*_*l*_, …, *λ*_*L*_ with *IF*_*j*_ means 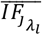. Then,

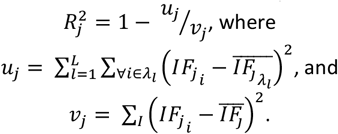

Once the 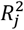 of *ϕ*_*j*_ is calculated on *ϕ*_*j*_(*X*_*j*_) → *y*_*j*_, one of the covariate columns of the *X*_*j*_ matrix is randomly permuted to form 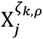, where *ς*_*k, ρ*_ denotes a random permutation *ρ* ∈ P of the covariate *x*_*k*_ column. 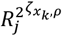 is then calculated on 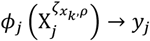. The importance of covariate *x*_*k*_ is then defined as the decrease in score:

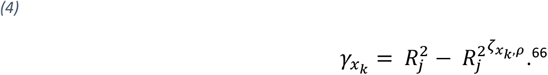

Although the significance criteria can be changed by the user, in the default settings of SPIT a candidate transcript is only labeled as DTU with the following condition:

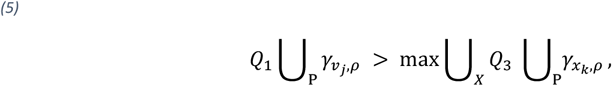

where *Q*_1_ and *Q*_3_ refer to the 1^st^ and 3^rd^ quartiles of the permutation importance scores, respectively. The number of permutations for the permutation importance test is a user-set parameter and defaults to 100.

### Parameter-fitting

SPIT has two main hyperparameters that affect its behavior: bandwidth (*h*) for KDE-fitting, and *κ* for *p*-value thresholding. The choice of bandwidth (*h*) directly determines the level of smoothing in the KDE function, with larger values of *h* leading to oversmoothed and smaller values leading to undersmoothed *IF* distributions.^68^ In contrast to the conventional interpretation of an optimal bandwidth, selecting an optimal bandwidth for SPIT does not require achieving the highest possible accuracy in representing the underlying *IF* histograms. This is due to the fact that overdispersion in RNA-Seq data can lead to overly erratic histograms, which may be identified as multimodal by traditional approaches. Rather, selecting high values of *h* allows us to reduce the risk of false discoveries by “oversmoothing” the input *IF* distributions and only detecting only the most significant partitions in the data.

Similar to the choice of bandwidth, the optimal *κ* value also depends on the level of dispersion present in the input dataset. Smaller values of *κ* lead to more stringent behavior by setting smaller *p*-value thresholds for detecting DTU events. To estimate the optimal values of *h* and *κ* for each dataset, SPIT implements a parameter-fitting process similar to cross-validation. This involves creating a set of experiments by introducing simulated DTU events into the input control group, following the same approach as used in the GTEx test experiments. Then, different combinations of *h* and *κ* values are evaluated based on their accuracy.

Given the set of case samples *I*_*case*_ and the set of control samples *I*_*control*_, we define a number (*n*_*e*_) of experiments, 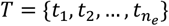. To simulate each of the parameter-fitting experiments:

1. Randomly divide *I*_*control*_ into two sets of equal size to create the simulation case and control groups, 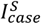 and 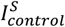, respectively.
2. Apply the SPIT pre-filter outlined above assuming the randomly assigned 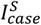 and 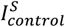. As with the GTEx test experiments, we skip step 6 of the pre-filtering process.
3. We repeat the steps 3-5 of the GTEx test experiment simulation on 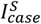 and 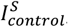, where the number of splicotypes introduced into 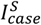 is a user-set parameter (*n*_*g*_, defaults to 5). For simple genetic disorders and experiments with small sample sizes, *n*_*g*_ can be set to 1 as a complex partition within the case group is either not expected or cannot be detected. The noise parameter *ϵ* can also be set by the user, and defaults to 0.05 as in the GTEx simulation.

In order to estimate the optimal values of *h* and *κ* (i.e. *h*^*^ and *κ*^*^) out of all combinations within user-set search ranges (with defaults 0.02 ≤ *h* ≤ 0.20 and *κ* ∈ {0.1, 0.2, …, 1}), we employ a leave-one-out cross-validation (LOOCV) approach on the simulated set of experiments, *T*. For each step *s* in *n*_*e*_ number of iterations:

1. Let *T*_(*s*)_ = *T \ t*_*s*_. We run SPIT on *T*_(*s*)_ with all (*h*_*i*_, *κ*_*j*_) | *h*_*i*_ ∈ {0.02, 0.03, …, 0.20}, *κ*_*j*_ ∈ {0.1, 0.2, …, 1} to yield estimated *F*-scores, 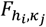.
2. Select 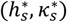 such that 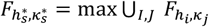.
3. Run SPIT on *t*_*s*_ with 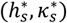 to get *F*.

After *n*_*e*_ iterations, we obtain a set of optimal hyperparameters and their corresponding *F*-scores: 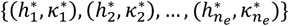and 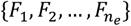. We select the hyperparameter values with the highest consensus among the iterations as our estimated optimal values (*h*^*^, *κ*^*^). The average *F*-score 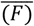 across all iterations can be interpreted as the overall *F*-score of the SPIT pipeline on the provided dataset, which can help determine if SPIT is an appropriate analysis tool for the dataset. In general, larger sample sizes of the control group (*n* ≥ 16) are expected to improve accuracy of SPIT test as the *U*-statistic is nearly normal with *n* = 8 vs. 8.^69^ Consequently, the parameter-fitting experiments are expected to reveal the best results with control group sizes ≥ 32.

For the parameter-fitting experiments in this work, we used the default search ranges with *n*_*e*_ = 10 and *n*_*g*_ = 5. (*h*^*^, *κ*^*^) were estimated as (0.09, 0.6) for the GTEx simulation experiments, and (0.06, 0.4) for the Lieber brain samples. Final 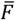 across 10 experiments were 0.911and 0.930, respectively.

SPIT’s parameter-fitting process can be time-consuming and computationally intensive, and it is an optional step. Running 10 experiments (*n*_*e* = 10) on a typical personal laptop can take up to 24 hours, however, multithreading is available through GNU parallel.^70^ Without parameter-fitting, the default values of (*h, κ*) are set to the estimated optimal (*h*^*^, *κ*^*^) for the GTEx dataset, (0.09, 0.6).

### Removing outlier effects and tie-correction

Assume that a global minimum was detected in the *IF* distribution of case samples in order to partition subgroups for an arbitrary transcript, and the left and right tails of the case and control groups were determined as *l*_*case*_, *r*_*case*_, *l*_*control*_, and *r*_*control*_.

We define a parameter *n*_*small*_, which defines the minimum size for subgroups that can be confidently detected and interpreted in the given dataset. If either or both of the sizes of *l*_*control*_ and *r*_*control*_ are smaller, they can be expanded to the right and to the left, respectively, until each contains at least *n*_*small*_ samples for comparison. Unlike the tails of the control group, *l*_*case*_ and *r*_*case*_ represent meaningful stratifications within the case group that may have biological implications. Therefore, the group sizes of both *l*_*case*_ and *r*_*case*_ need to be at least *n*_*small*_. Otherwise, the stratification is considered unreliable due to potential influence of outliers. In such cases a Mann-Whitney *U* test is conducted between the entire groups of *I*_*case*_ and *I*_*control*_.

Additionally, in order to reduce the impact of insignificant differences between *IF* values in the Mann-Whitney *U* test, SPIT rounds all *IF* values to two decimal points.^71^ The normal approximation for the *U*-statistic^69^ is used for tie-correction for group sizes larger than 8. Although SPIT works well with smaller sample sizes (*n* ≥ 12) for simple genetic architectures, it requires *n* ≥ 24 samples for each group for the normal approximation to be reliable in SPIT-Test module. Exact *U*-statistic *p*-values are computed for group sizes smaller than 8 when there are no ties.

## Data availability

The “Swimming Downstream” dataset is uploaded to Zenodo by Love *et al*.:

Quantification files: https://zenodo.org/record/1291522

Scripts and simulation data: https://zenodo.org/record/1410443

All 20 of the GTEx simulation experiments are uploaded to Zenodo: https://zenodo.org/record/8128846

Quantification files and phenotype information for the GTEx samples used in the detection of tissue-dependent DTU events are uploaded to Zenodo: https://zenodo.org/record/8128945

The RNA-Seq data used in the schizophrenia analysis are made available by the Lieber Institute for Brain Development at http://eqtl.brainseq.org/phase2/.

## Acknowledgements

This work was supported in part by the U.S. National Institutes of Health under grants R01-MH123567 and R01-HG006677.

## Notes

### Competing Interest Statement

The authors have declared no competing interest.

