## Supplementary Methods for "Detecting differential transcript usage in complex diseases with SPIT"

**Sashimi plots for tissue-dependent DTU events**

To obtain the necessary data, we aggregated the read alignments from all samples in each tissue using TieBrush^1^ and used its module TieCov to extract base-pair and junction coverages.

To manually validate the presence of differentially expressed signals between transcripts at a locus, we constructed sashimi plots for each gene in the evaluation.^1, 2^ These plots, shown for each gene in Figure 3, depict the coverage from each tissue tested for that specific locus. All coverage values obtained using TieCov were normalized using the following formula:

$$\left( \frac{C_{i}}{\sum_{j=0}^{N} C_{j}} \right)\cdot{10}^{6}$$

where $C_{i}$ represents the coverage at a given position being normalized, and $N$ is the length of the locus.

To assess differences in the transcriptional landscapes between tissues at each locus, we calculated the change in coverage compared to the average across all GTEx samples (Δ). In Figure 3, the Δ track represents the results obtained by subtracting the normalized coverage values of each tissue from the normalized coverage of the entire GTEx dataset.

In order to run SPIT, all samples were quantified using Salmon^3^ with CHESS 3^4^ as reference annotation.

**Filtered-CPM threshold**

It is worth noting that although $IF_{i,j}$ values are not measures of gene expression, they may still be affected negatively by extremely low gene expression values. For an arbitrary transcript $j$ of gene $g$, let samples $a$ and $b$ both have $g_{c}=10$, and $t_{a, j}=2$, $t_{b, j}=6$, respectively. As a result we get $IF_{a, j}=0.2$ and $IF_{b, j}=0.6$, which seem to indicate a significant DTU while in reality a difference of 4 in read count is negligible. Therefore, in order to avoid disproportionally inflated differences in $IF_{j}$ values, SPIT has an optional Filtered-Counts per million (CPM) threshold which, for a transcript $j$of gene $g$, only considers the samples with CPM $\geq10$ for $g$ in the Mann-Whitney $U$ test. CPM values for this threshold are calculated on the selected subset of genes that pass the pre-filtering steps above, assigning the total count of these genes as the library size for each sample. This threshold is only used with the real RNA-Seq datasets analyzed in this paper, excluding all simulated experiments.

**Flagging DTU genes based on likelihood scores**

The KDE-fitting step of SPIT estimates a smoothed distribution for the $IF$ values of each transcript in the case and control groups, which can be exploited to further evaluate candidate DTU events. For an arbitrary DTU event in transcript $j$ between case group $I_{case}$ and control group $I_{control}$, let the estimated kernel densities for $I_{case}$ and $I_{control}$ $IF$s be ${K'}_{case}$ and $K_{control}^{'}$, respectively.

In addition to the Mann-Whitney $U$ statistic between the $IF$ distributions of $I_{case}$ and $I_{control}$, we also calculate the likelihood scores of all $\bigcup_{i\in I_{case}} IF_{i,j}$ using density function $K_{control}^{'}$, denoted as $L_{j}$. This gives us a measure of the probability of observing the $IF$ values of the case group given the $IF$ distribution of the control samples. Upon collecting the likelihood scores of all transcripts $\bigcup_{J} L_{j}$, we label outlier transcripts using a median absolute deviation (MAD) test with the conventional threshold of $3.5$.^5^ As with the $U$-statistic $p$-values, presence of subgroups within the case samples results in two separate likelihood scores for a single transcript, in which case the smallest likelihood score gets assigned to the transcript. We assign a significance flag to any candidate DTU gene that has at least one identified outlier transcript.

**Quantification of the GTEx heart (left ventricle) samples**

Transcripts were quantified using Salmon^3^, using the entire genome GRCh38.p14 as a decoy sequence and the reference annotation RefSeq (release 110).^6^ TPM values computed with Salmon were scaled up to library size using the “scaledTPM” conversion from *tximport^7^*. All downstream analyses used scaled read counts as the unit of expression measurement.

**Assessment and quantification of Lieber Brain samples**

The sequencing quality of all brain RNA-Seq samples were assessed with FastQC^8^ and MultiQC^9^, and outlier samples were excluded from the analysis. Samples with post-mortem intervals of $\geq60$ hours were also excluded. Salmon^3^ was used to quantify all transcripts in the reference annotation CHESS 3^4^ using the entire GRCh38 genome as a decoy sequence. As with the GTEx samples, TPM values computed with Salmon were scaled up to library size using the “scaledTPM” conversion from *tximport*.^7^

As an extra quality control measure, we removed samples with a high proportion of genes exhibiting low expression. To do so, we first calculated the number of genes within each sample with Filtered-CPM ≤10. We then applied the median absolute deviation (MAD) test with a cutoff of $3.5$ to remove samples with a significantly higher number of low count genes. The entire pipeline is then rerun on the selected samples, including the pre-filtering steps.

**Use of Large Language Models (LLMs)**

ChatGPT^10^ has been used for refinement and polishing of text in this paper.
