## Supplementary Figures for "Detecting differential transcript usage in complex diseases with SPIT"

**Supplementary Figure 1**

*Experiment 2*

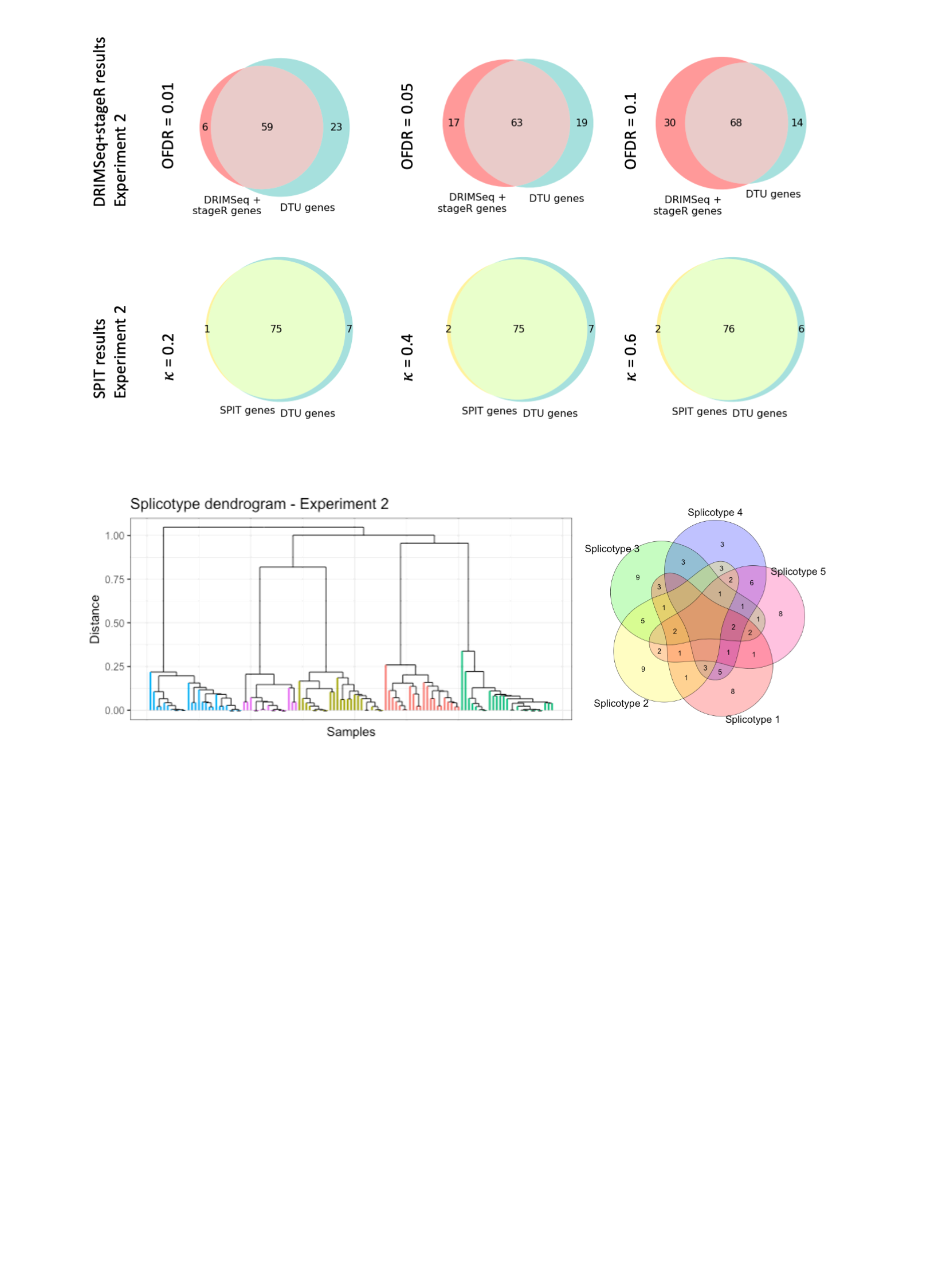

*Experiment 3*

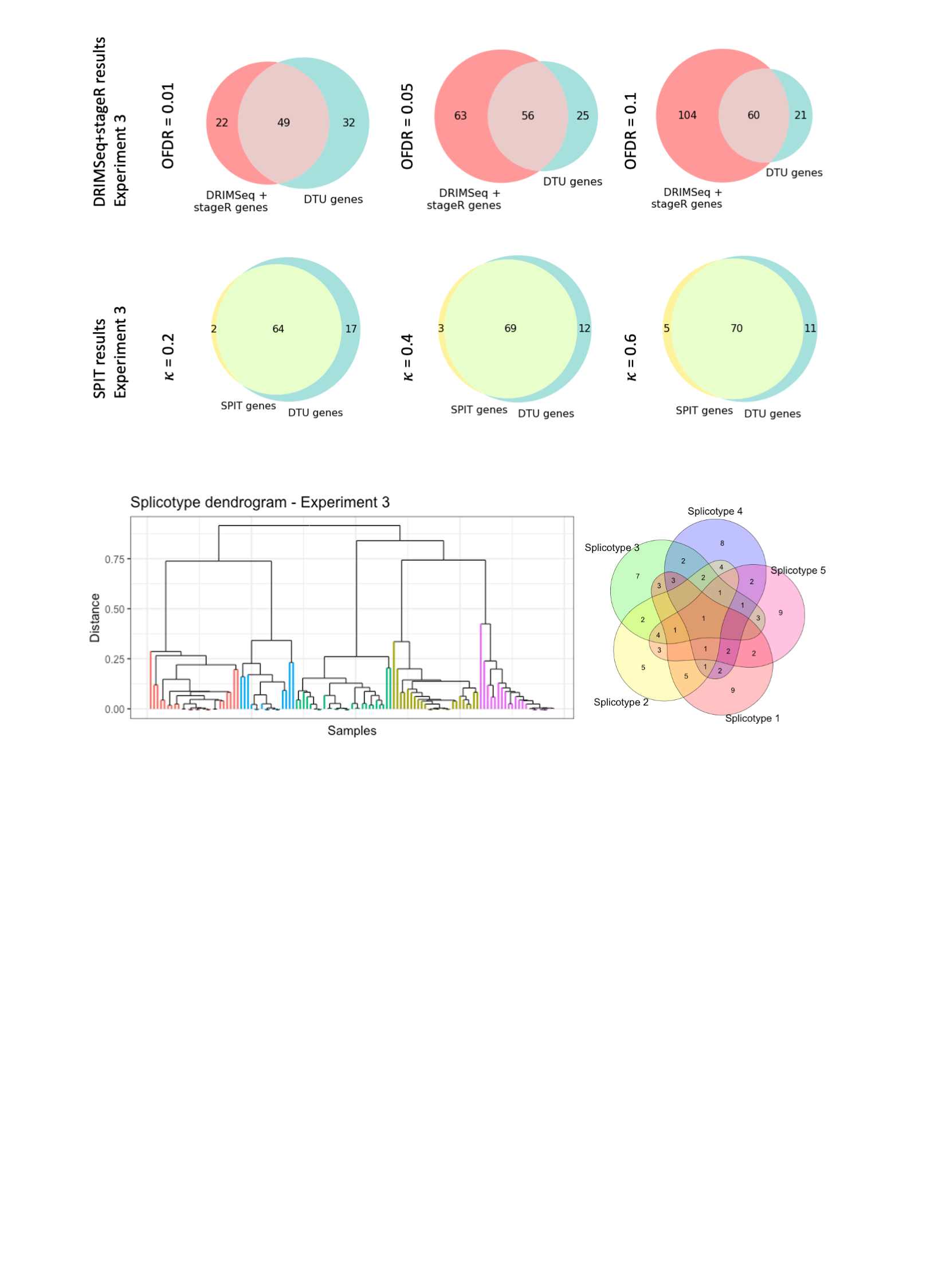

*Experiment 4*

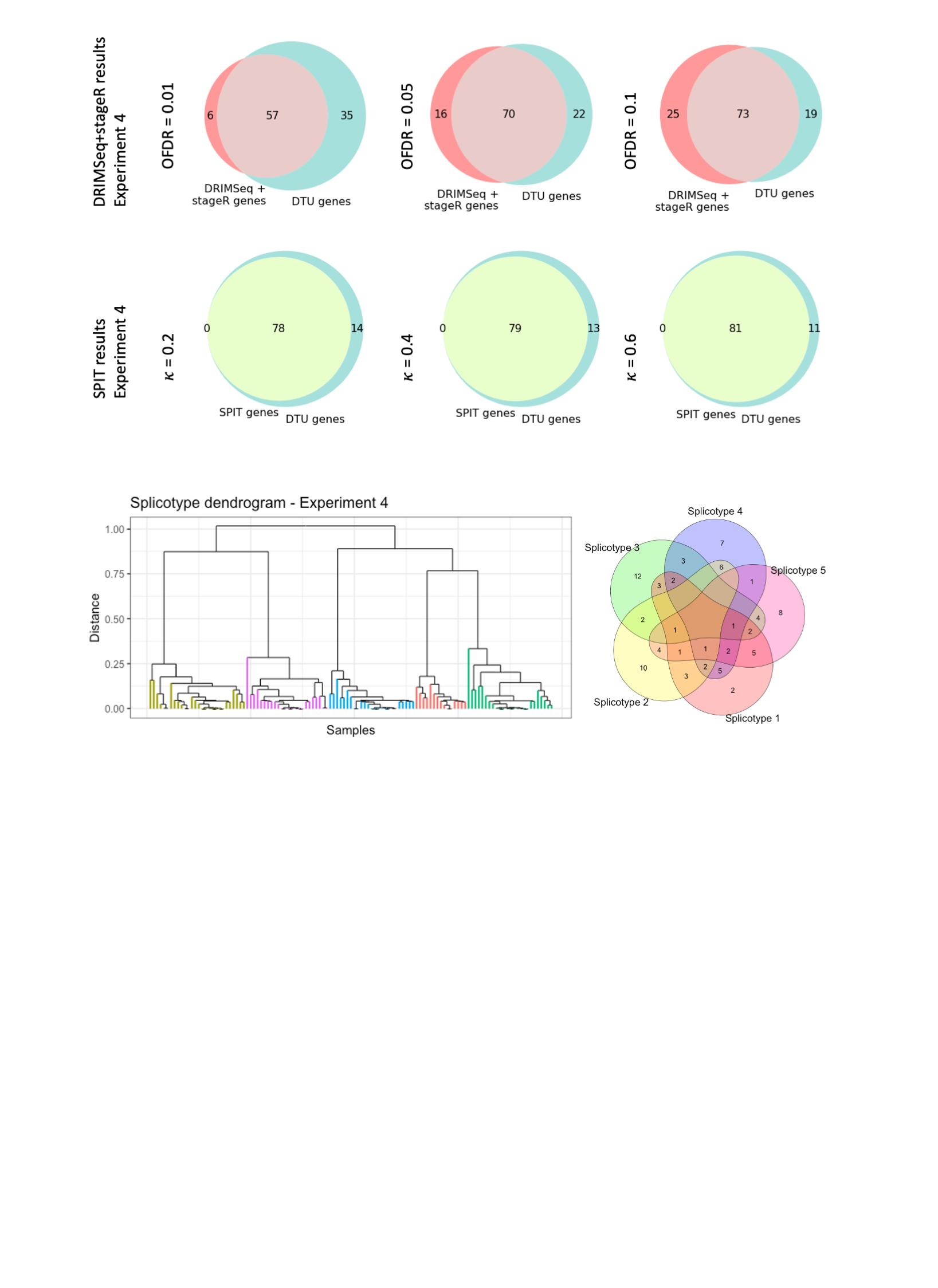

*Experiment 5*

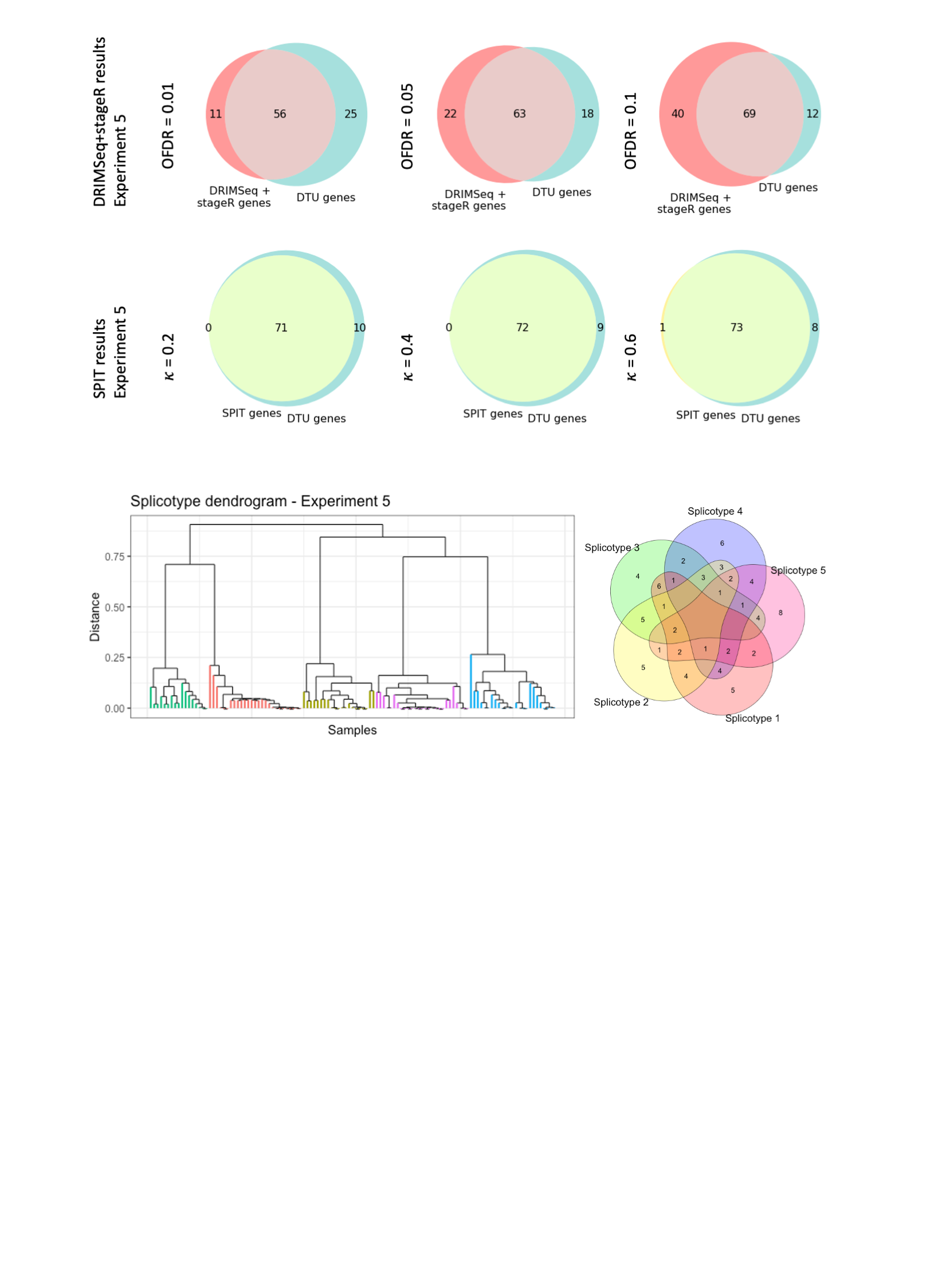

*Experiment 6*

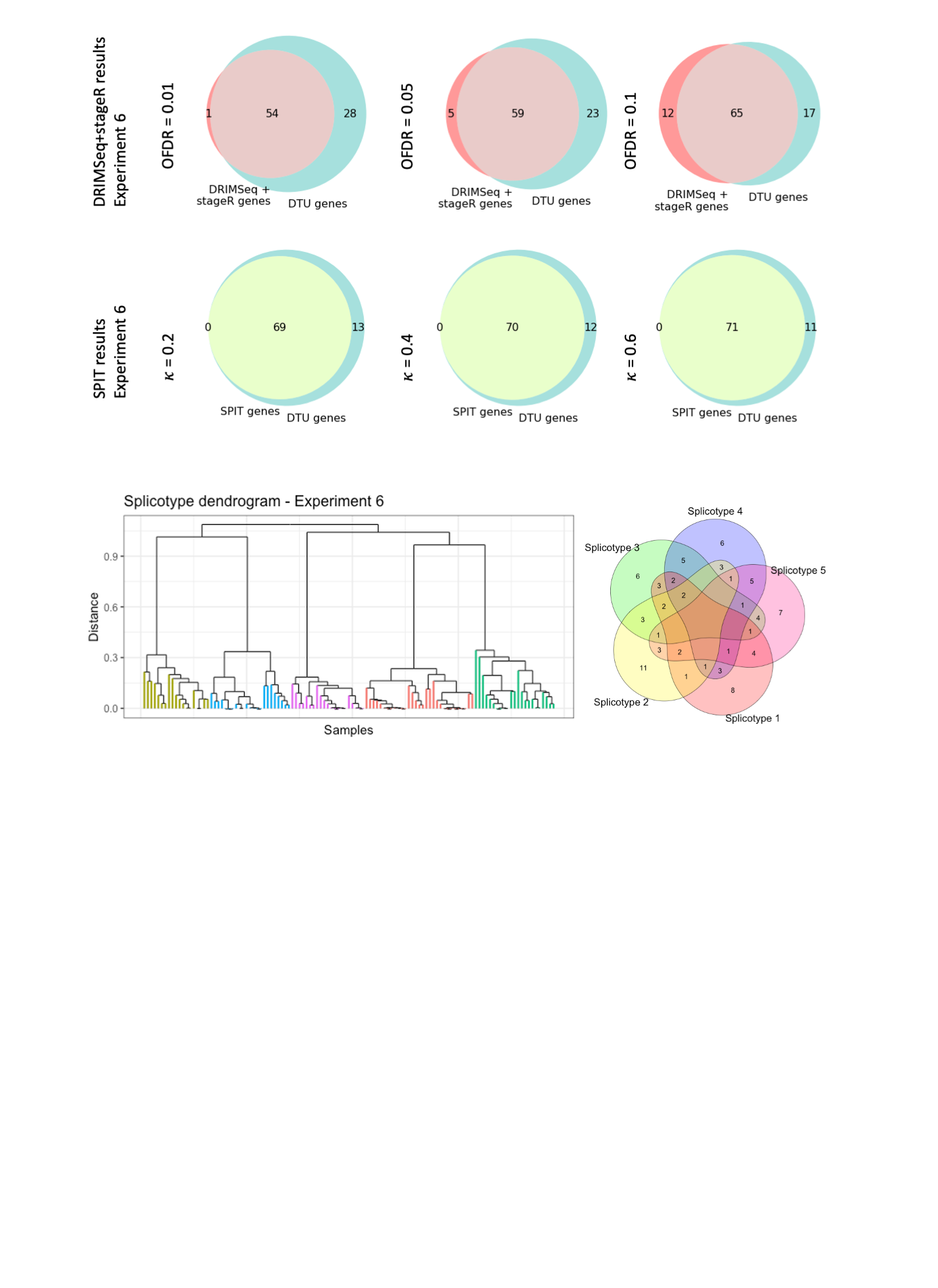

*Experiment 7*

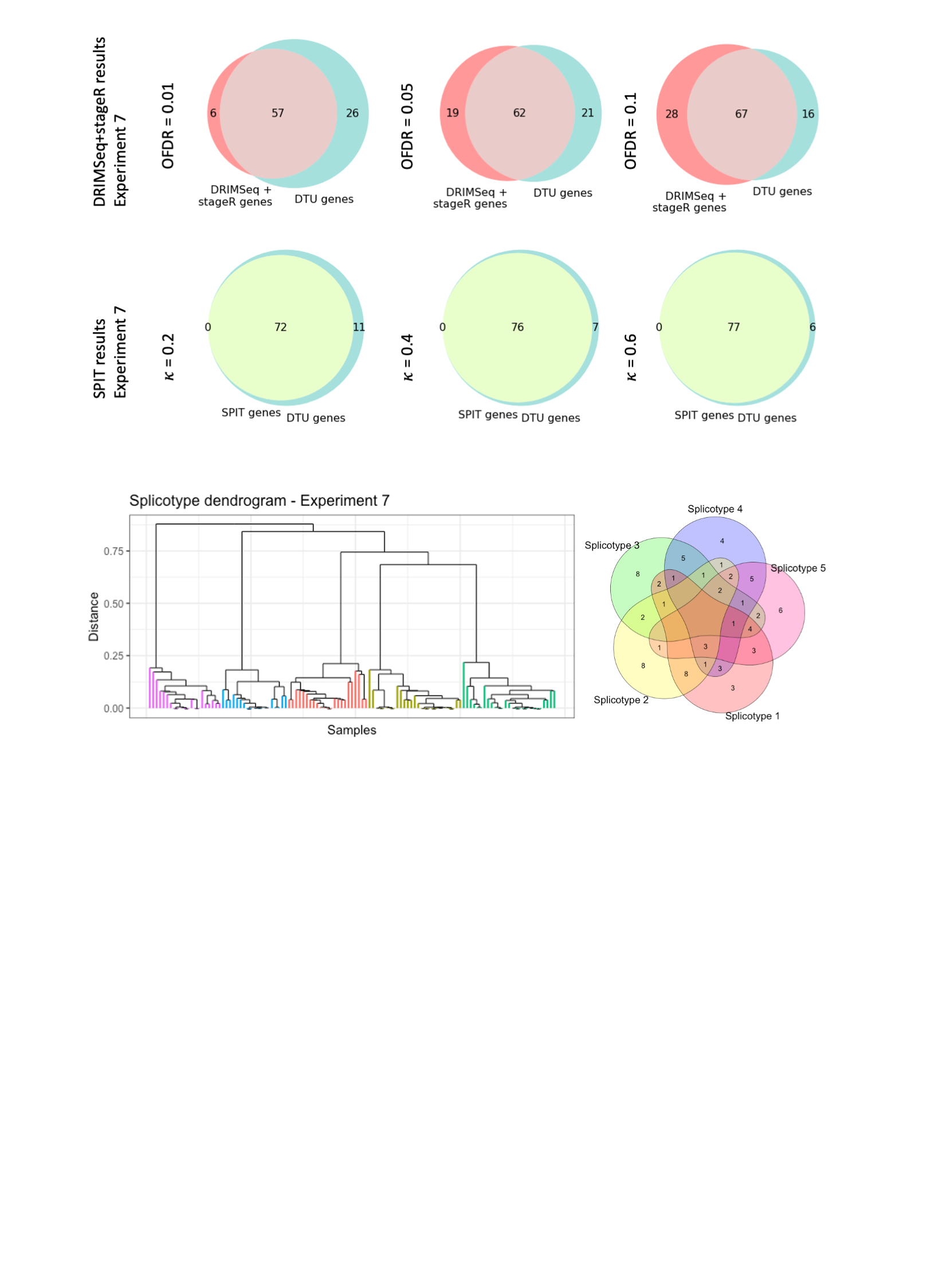

*Experiment 8*

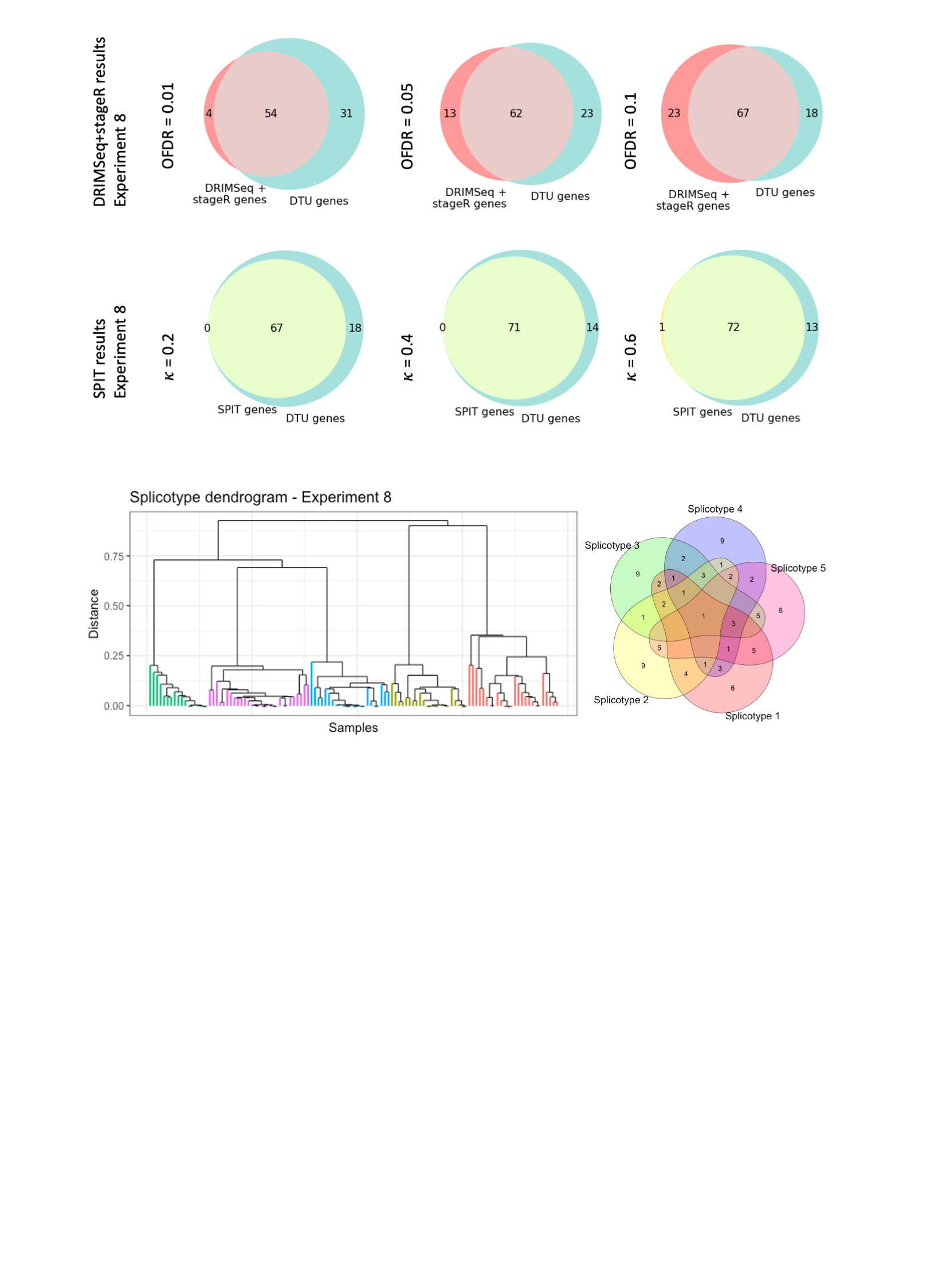

*Experiment 9*

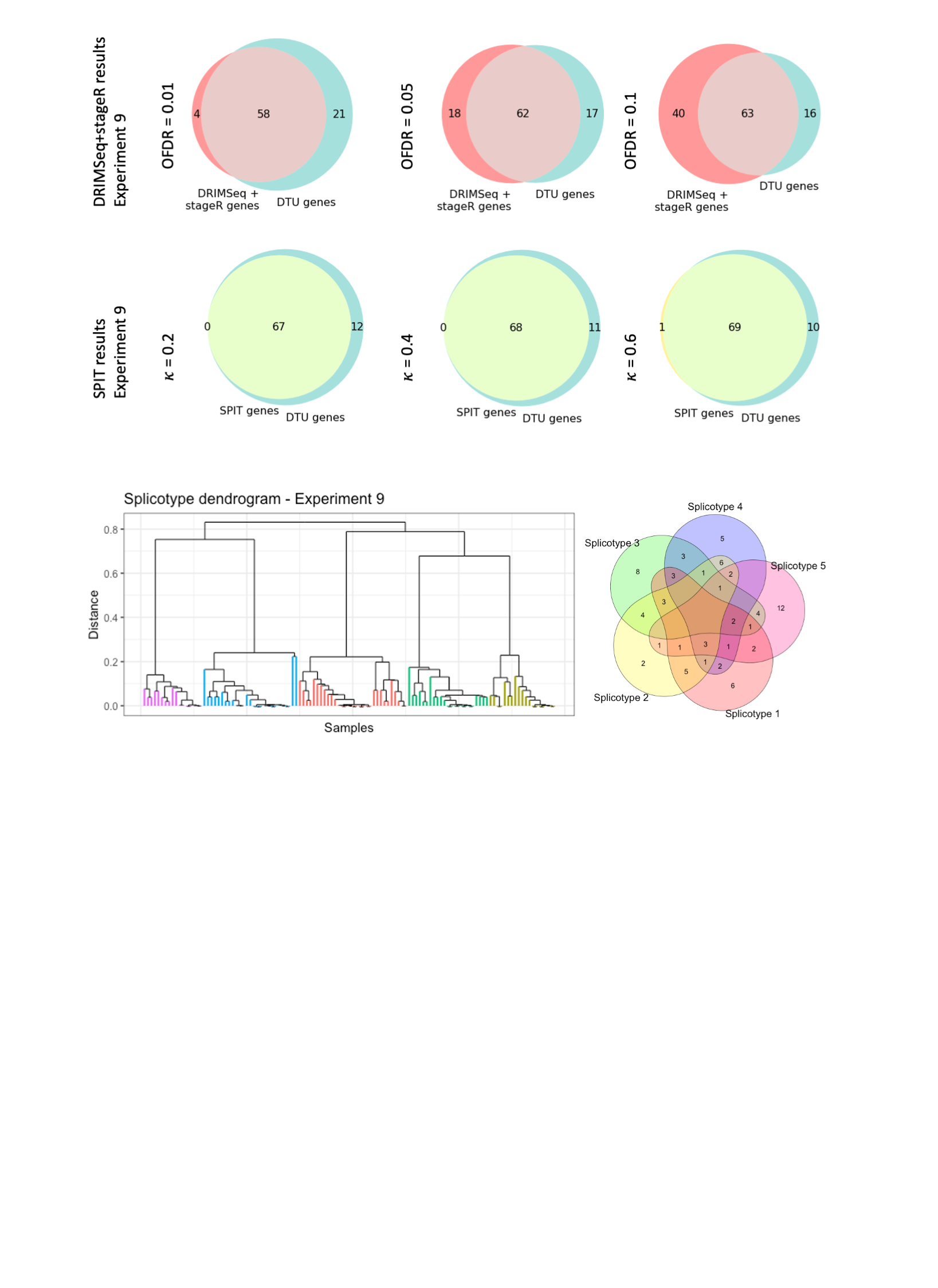

*Experiment 10*

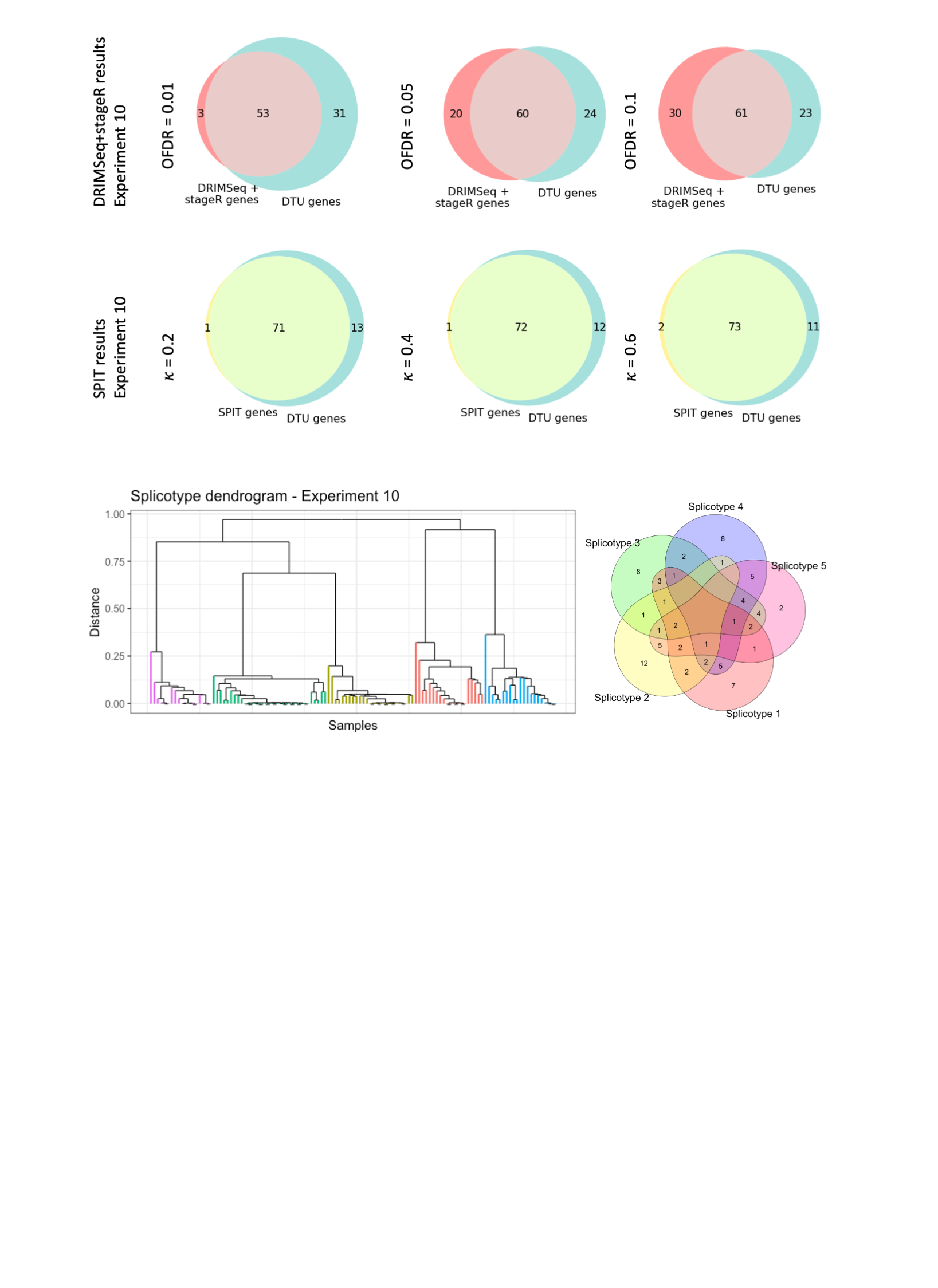

*Experiment 11*

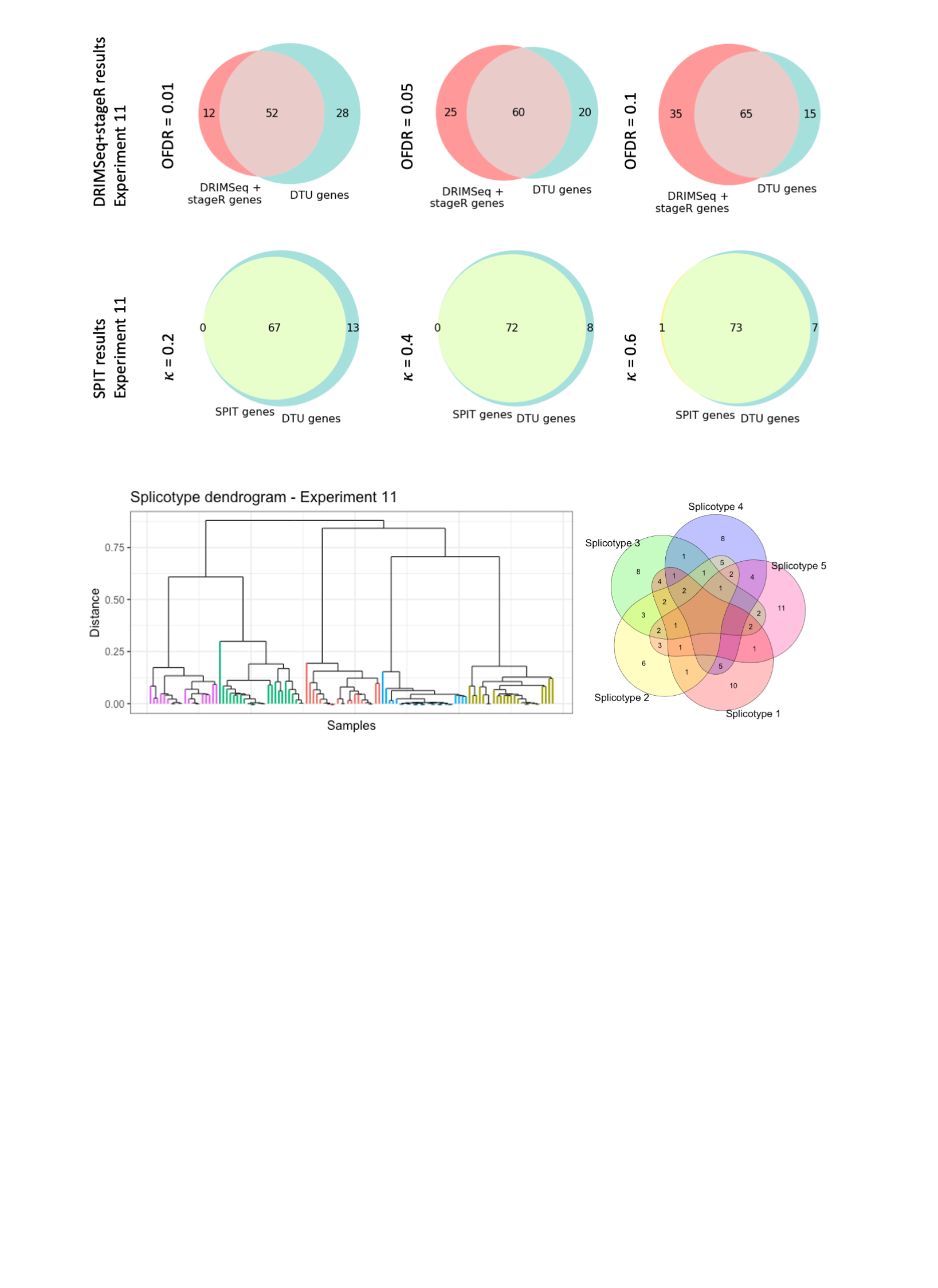

*Experiment 12*

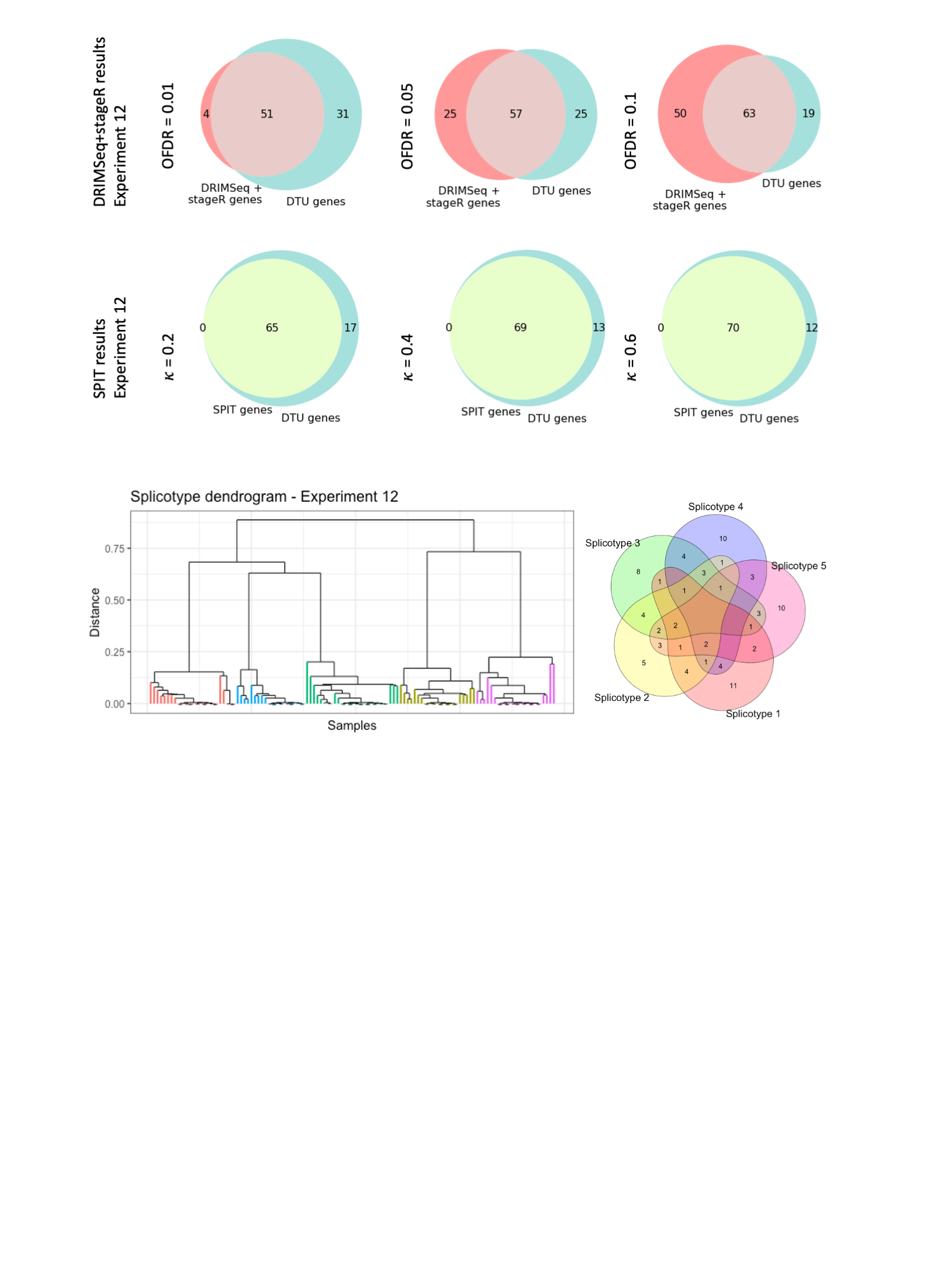

*Experiment 13*

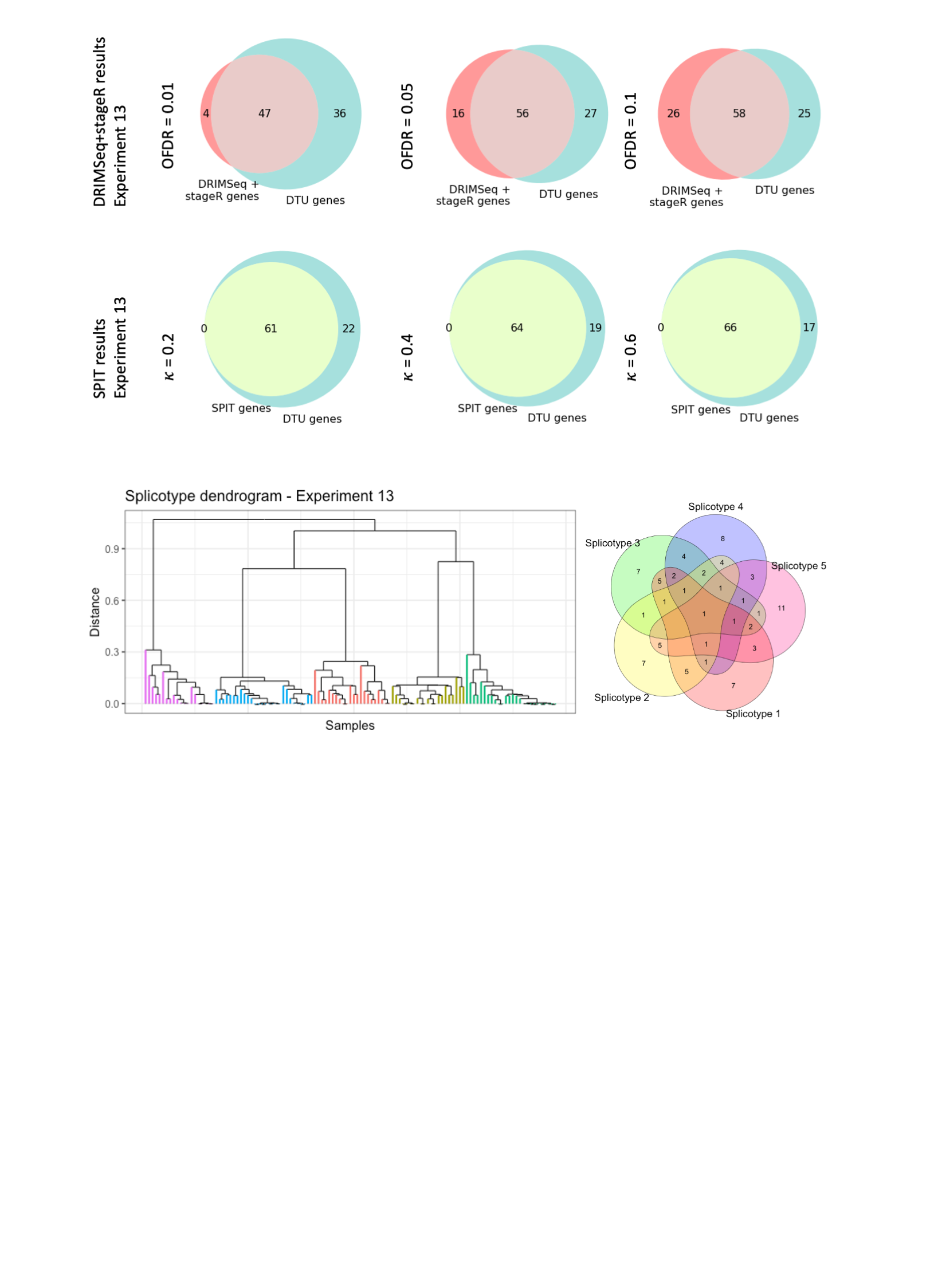

*Experiment 14*

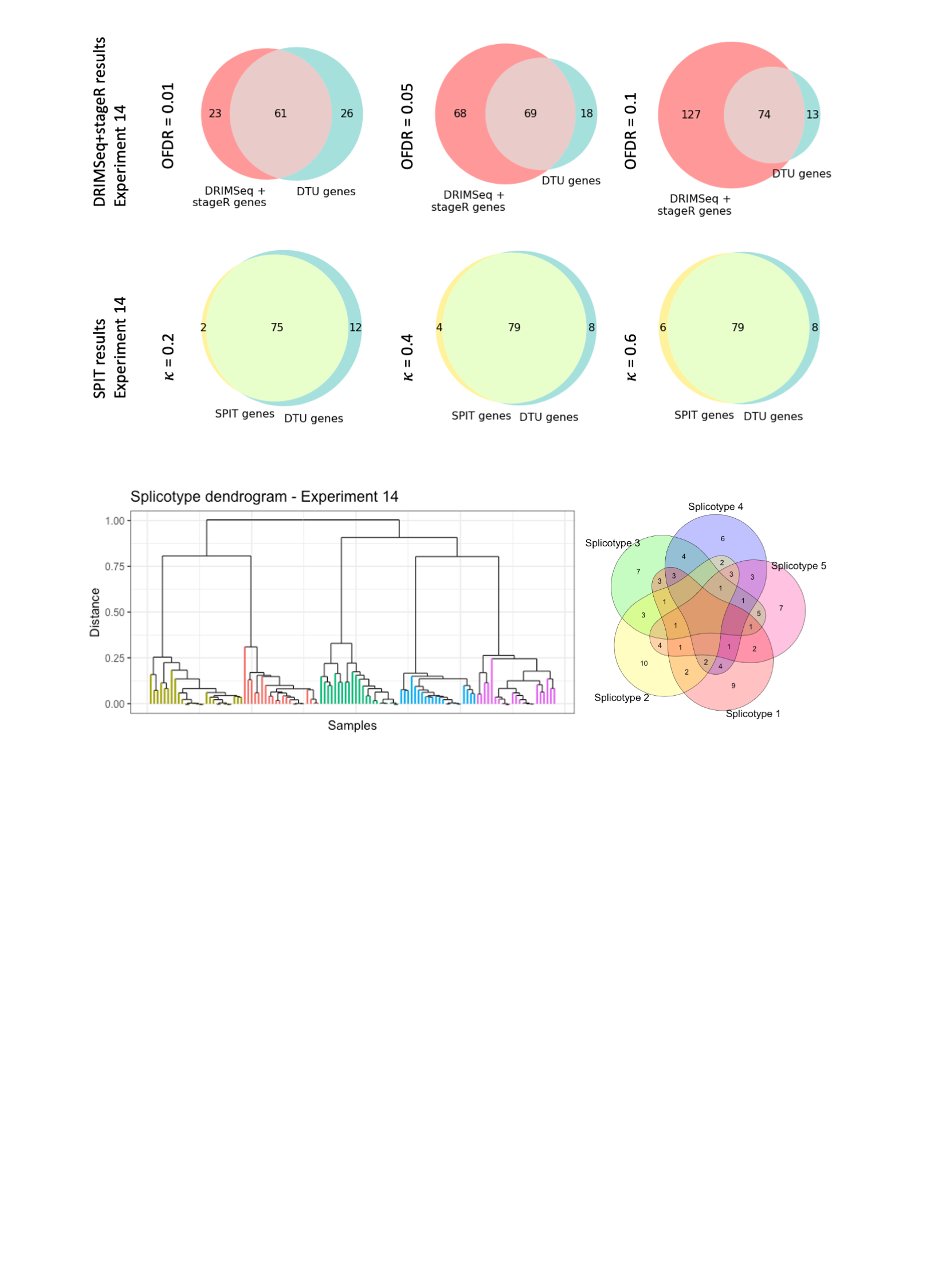

*Experiment 15*

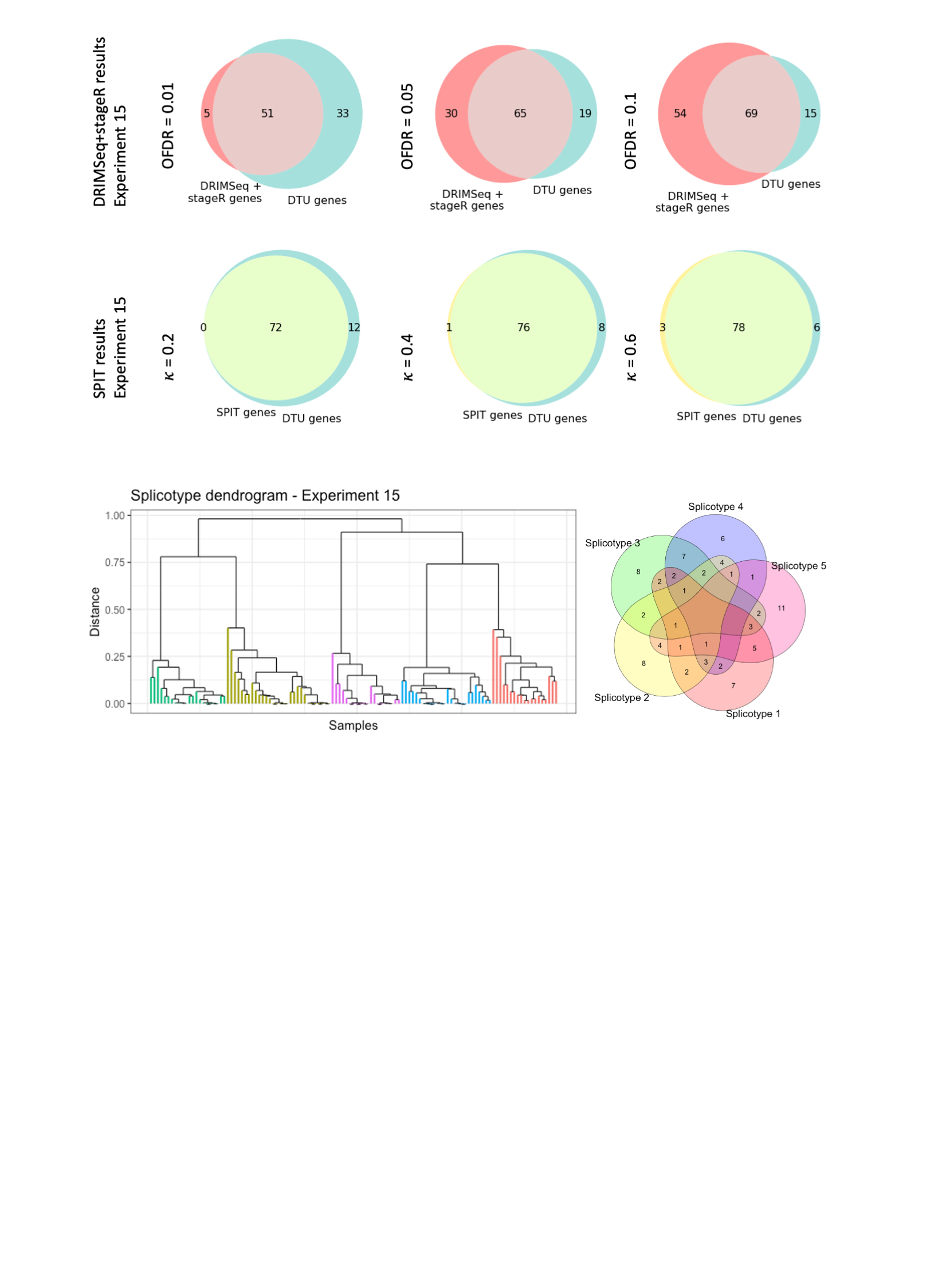

*Experiment 16*

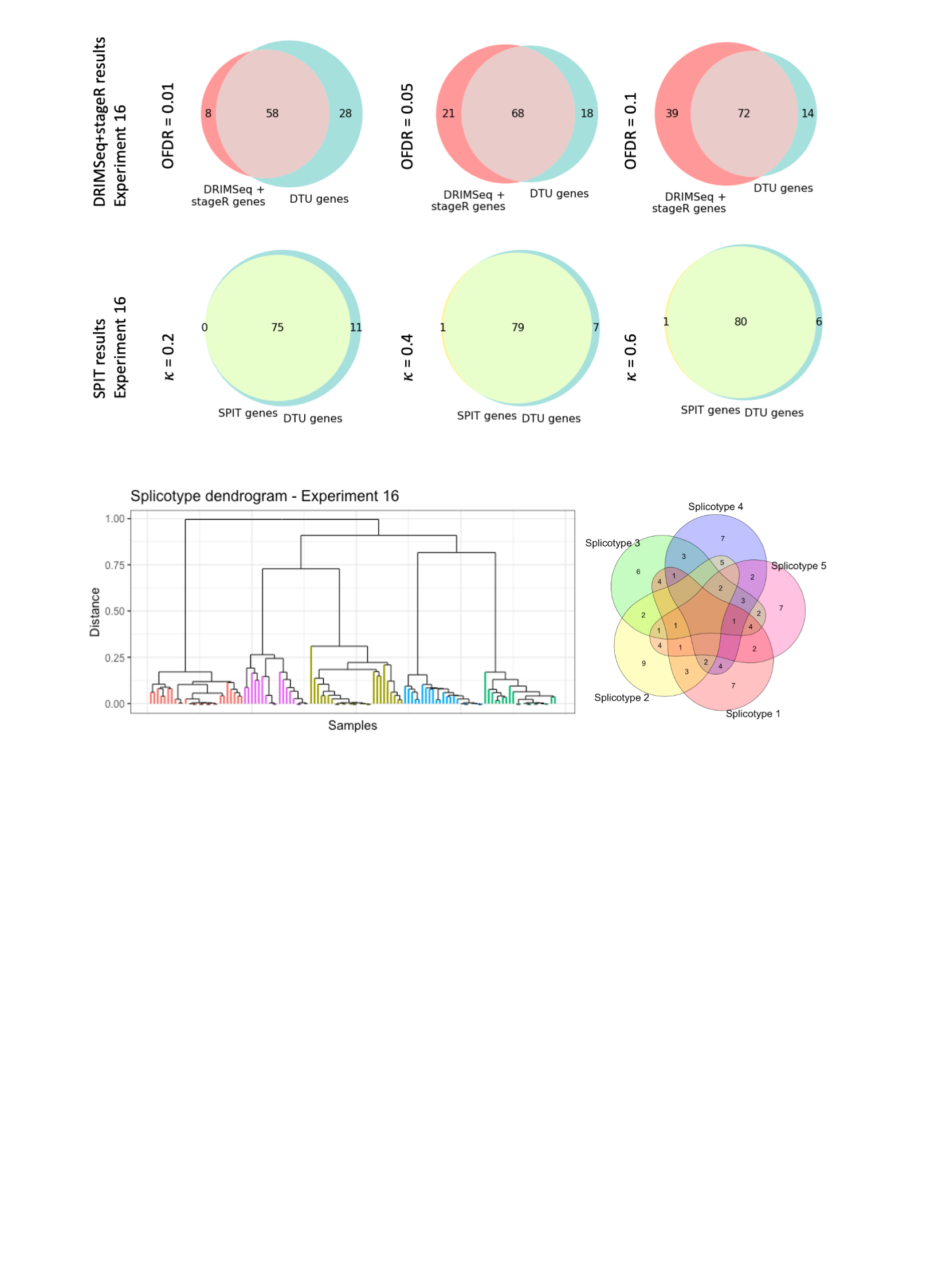

*Experiment 17*

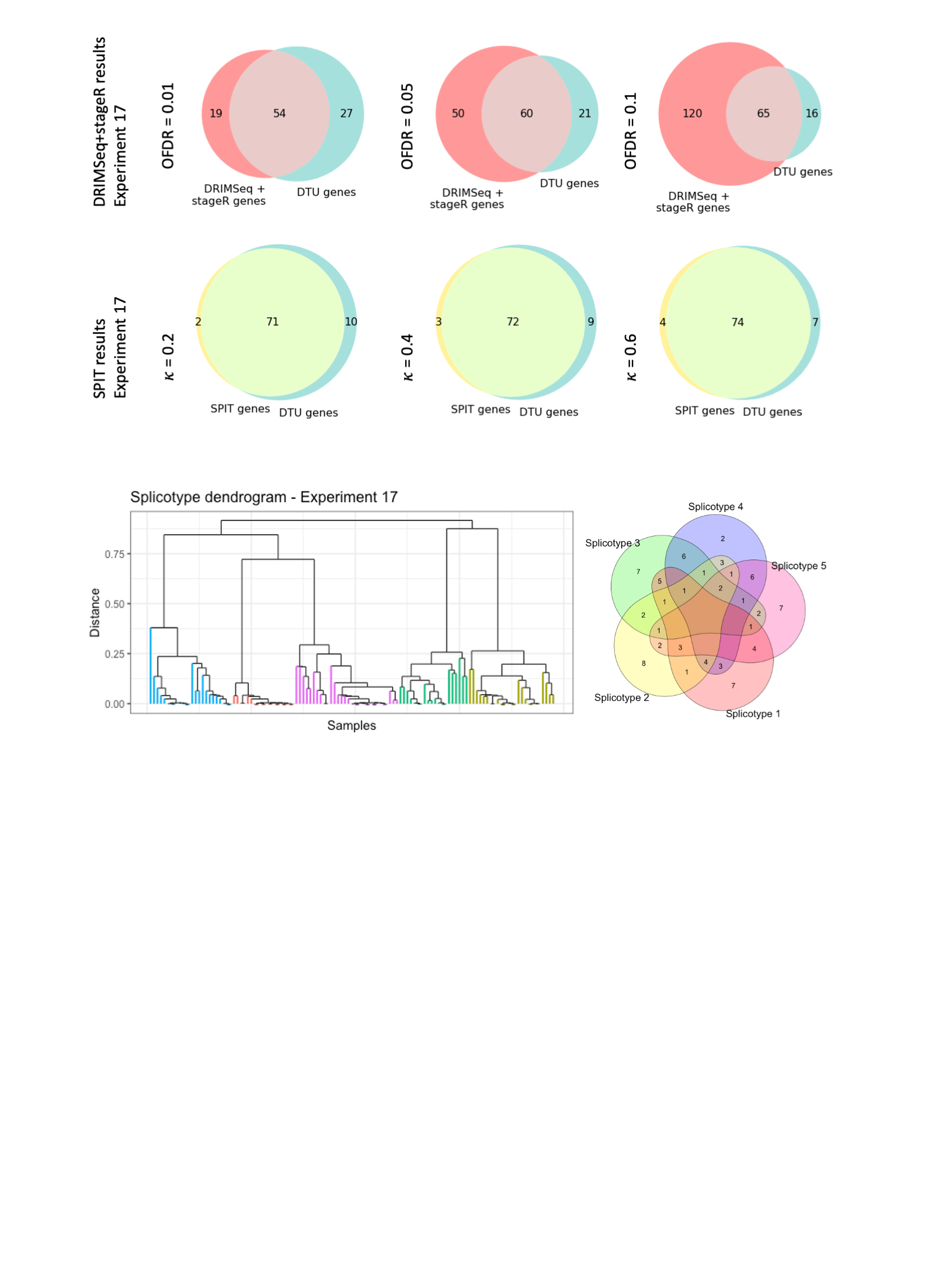

*Experiment 18*

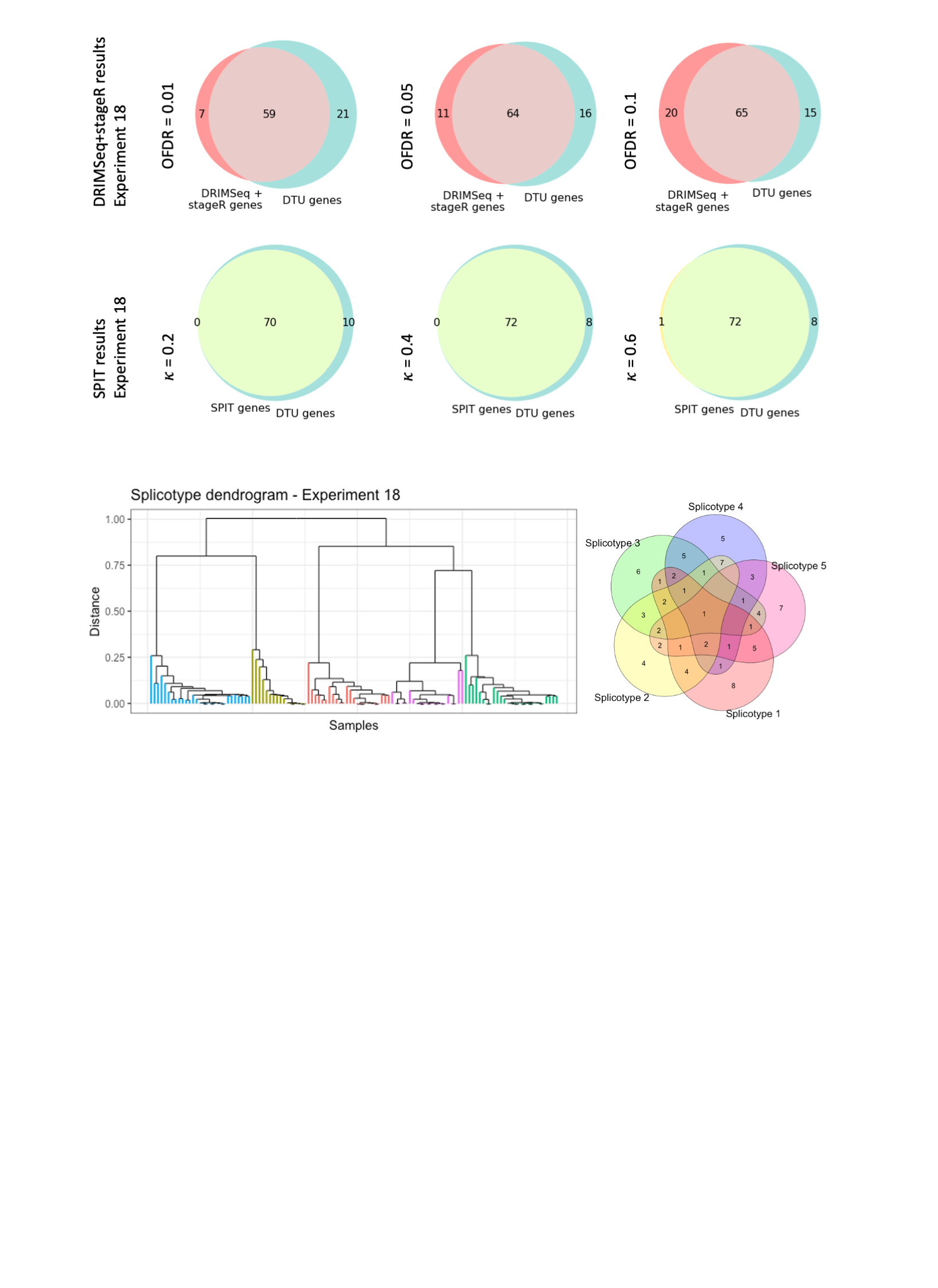

*Experiment 19*

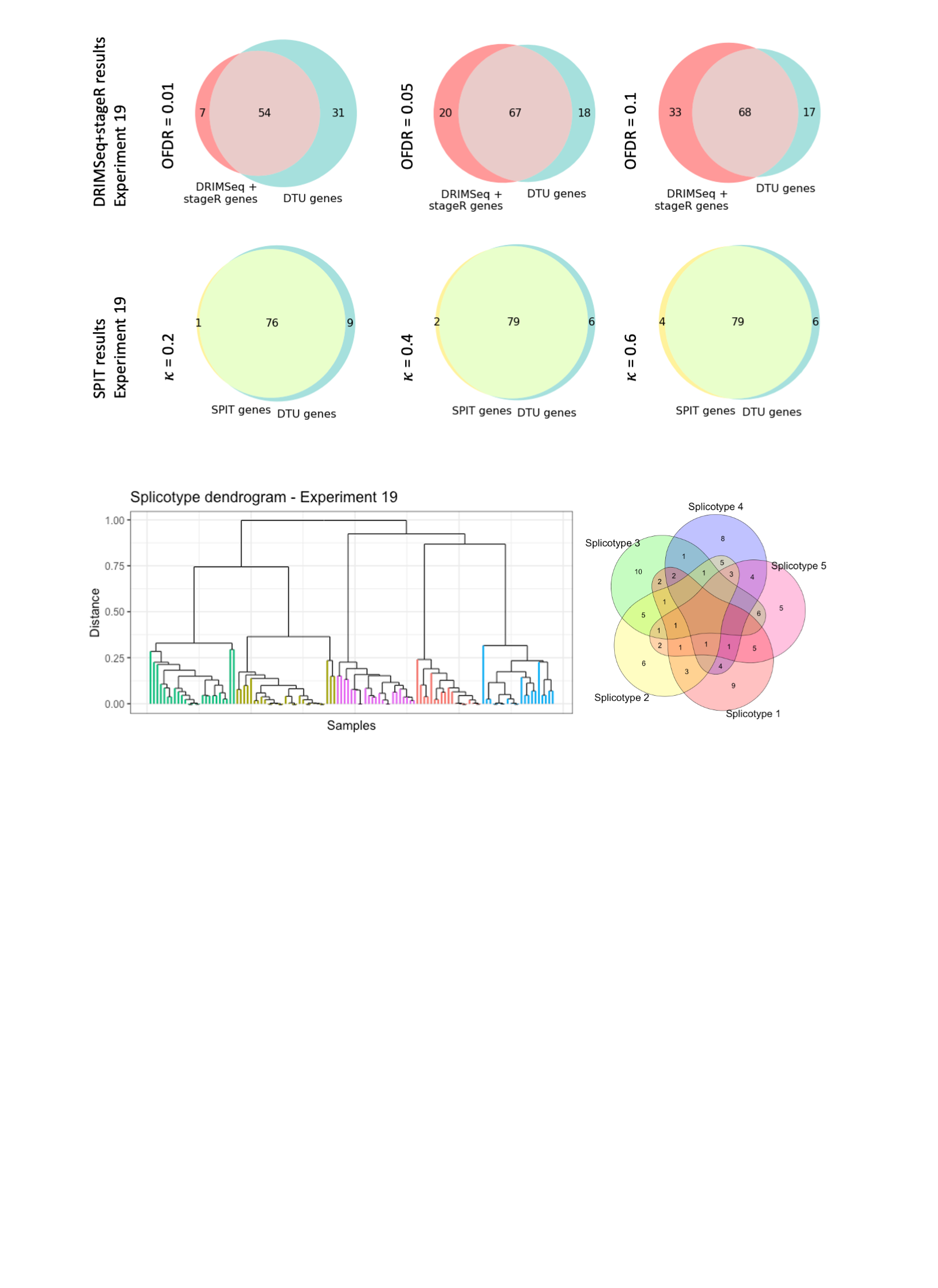

*Experiment 20*

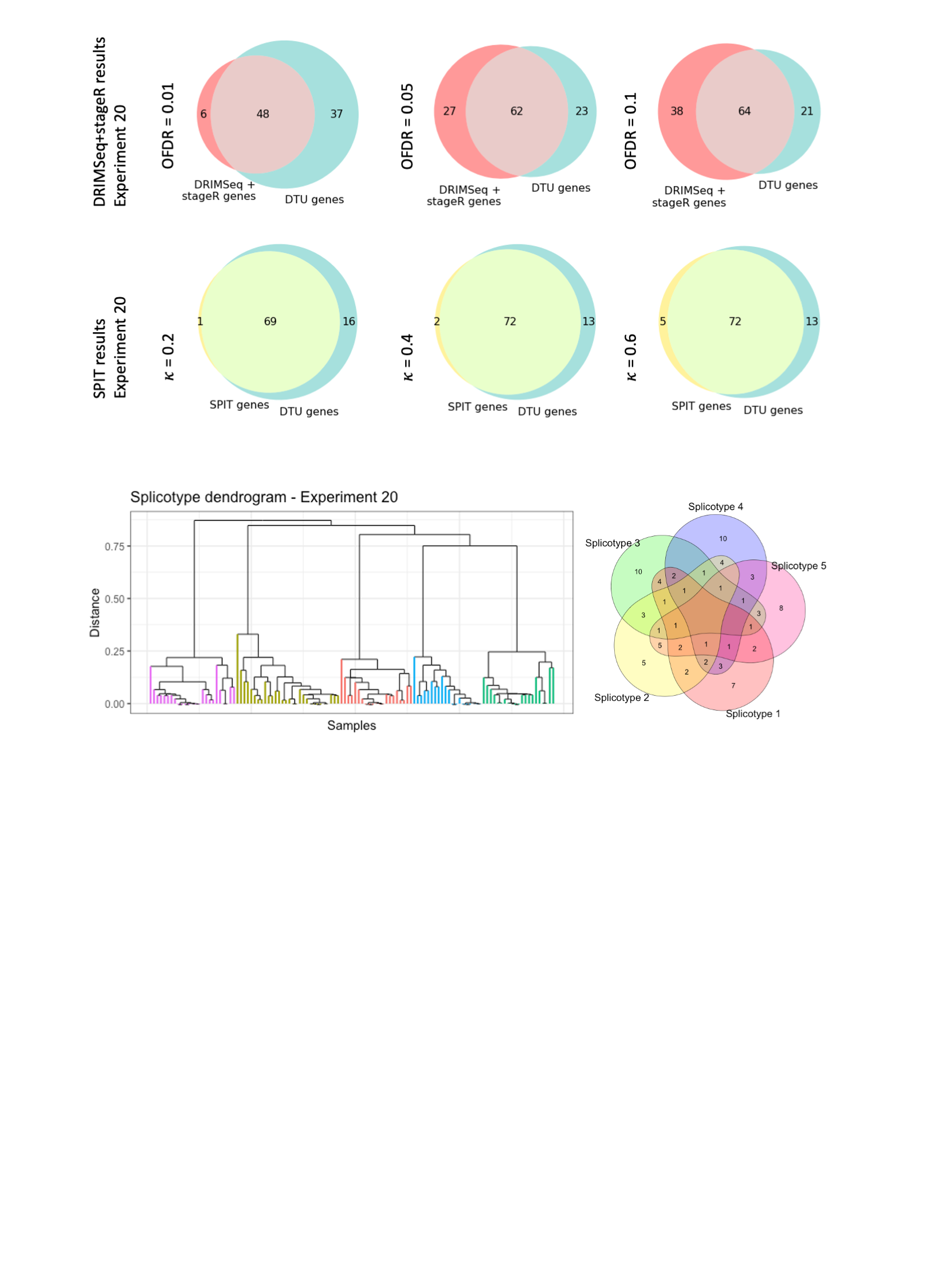

**Supplementary Figure 1:** Overlap of DRIMSeq+stageR pipeline and SPIT results with simulated DTU genes in experiments 2-19. The DTU event sharing Venn diagram and the corresponding final subcluster dendrogram based on the SPIT DTU matrix for experiments 2-19. The subclusters are color coded based on their distinct sets of simulated DTU events (splicotypes).

**Supplementary Figure 2**

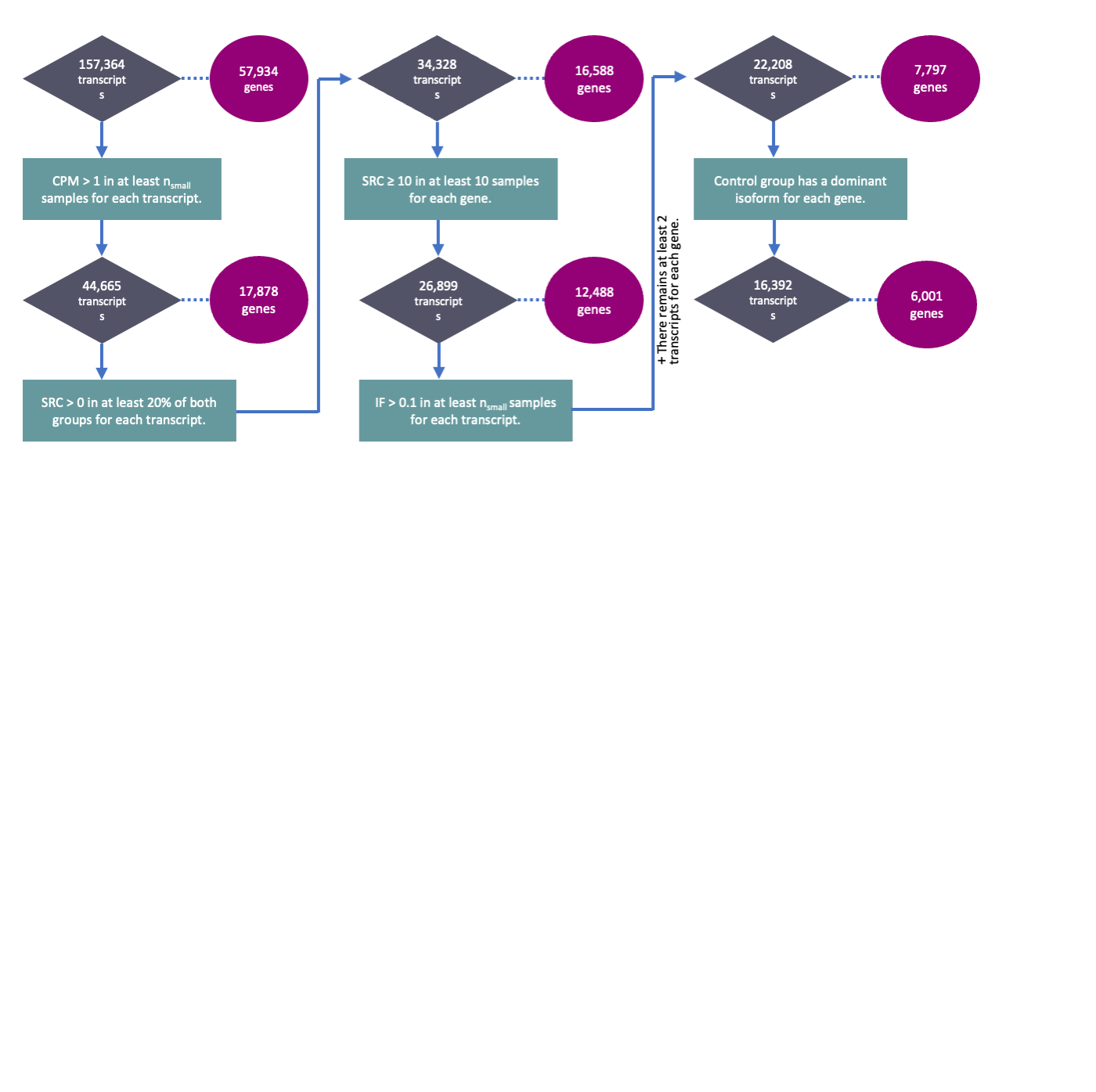

**Supplementary Figure 2:** Numbers of remaining transcripts and genes after each step of SPIT’s pre-filtering process applied on the Lieber brain samples with the default parameters.

**Supplementary Figure 3**

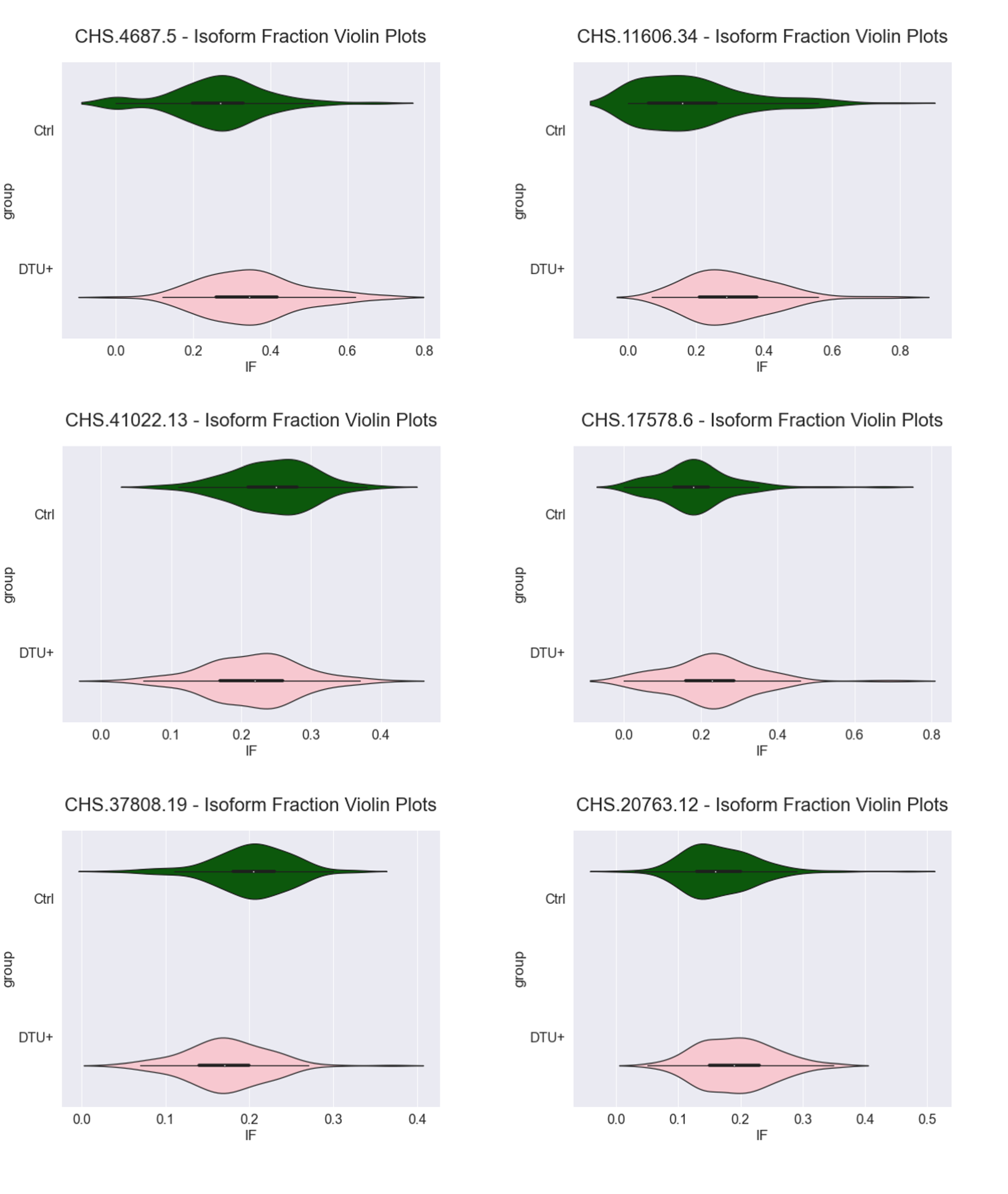

**Supplementary Figure 3:** Violin plots of the isoform fractions for the 6 transcripts of candidate DTU events in schizophrenia analysis.
