## Supplementary Table 1 for "Detecting differential transcript usage in complex diseases with SPIT"

| id | rin | condition | sex | race | age | batch |
|---|---|---|---|---|---|---|
| R12195 | 8.4 | 1 | M | AA | 62.61 | 2 |
| R12198 | 8.5 | 1 | M | CAUC | 29.98 | 2 |
| R12199 | 8.8 | 1 | M | CAUC | 65.26 | 2 |
| R12200 | 7 | 1 | M | AA | 32.35 | 2 |
| R12246 | 8.6 | 1 | M | AA | 52.18 | 2 |
| R12258 | 8.6 | 0 | F | CAUC | 69.07 | 2 |
| R12259 | 5.4 | 0 | M | CAUC | 48.1 | 2 |
| R12260 | 7.3 | 0 | M | AA | 50.28 | 2 |
| R12263 | 8.1 | 0 | F | AA | 47.01 | 2 |
| R12264 | 8.4 | 0 | M | CAUC | 49.26 | 2 |
| R12265 | 6.3 | 0 | F | AA | 35.21 | 2 |
| R12266 | 8.5 | 0 | M | CAUC | 54.43 | 2 |
| R12276 | 5.9 | 1 | M | AA | 75.22 | 2 |
| R12277 | 5.4 | 0 | F | AA | 50 | 2 |
| R12278 | 8.3 | 0 | M | AA | 24.79 | 2 |
| R12279 | 8.1 | 0 | M | AA | 58.48 | 2 |
| R12280 | 8.4 | 0 | M | AA | 22.09 | 2 |
| R12281 | 7.9 | 1 | F | AA | 60.75 | 2 |
| R12282 | 8.1 | 1 | M | AA | 47.35 | 2 |
| R12283 | 8.5 | 1 | M | AA | 35.82 | 2 |
| R12284 | 8.1 | 1 | M | CAUC | 25.59 | 2 |
| R12285 | 6.3 | 0 | F | AA | 58 | 2 |
| R12286 | 7.5 | 0 | M | AA | 40.39 | 2 |
| R12287 | 8 | 0 | F | AA | 19.69 | 2 |
| R12289 | 7.8 | 1 | F | AA | 61.93 | 2 |
| R12290 | 8.4 | 0 | M | AA | 45.91 | 2 |
| R12291 | 7.5 | 0 | M | AA | 18.05 | 2 |
| R12293 | 8.3 | 1 | F | AA | 62.63 | 2 |
| R12295 | 8.4 | 0 | M | AA | 41.55 | 2 |
| R12296 | 8.3 | 1 | M | AA | 38.72 | 2 |
| R12297 | 8.5 | 1 | M | CAUC | 43.38 | 2 |
| R12298 | 7.1 | 0 | M | AA | 36.95 | 2 |
| R12299 | 7.9 | 0 | F | AA | 30.48 | 2 |
| R12300 | 6.9 | 0 | F | AA | 52.57 | 2 |
| R12301 | 7.1 | 0 | M | AA | 57.89 | 2 |
| R12302 | 6.1 | 0 | M | AA | 57.1 | 2 |
| R12303 | 5.1 | 0 | F | AA | 48.42 | 2 |
| R12304 | 7.9 | 1 | M | AA | 63.18 | 2 |
| R12305 | 7.6 | 0 | M | CAUC | 63 | 2 |
| R12306 | 6.3 | 1 | F | AA | 41.59 | 2 |
| R12307 | 8.2 | 1 | F | AA | 49.65 | 2 |
| R12308 | 7.7 | 0 | M | CAUC | 57.6 | 2 |
| R12309 | 7.4 | 0 | F | AA | 57.59 | 2 |
| R12310 | 6.6 | 0 | F | AA | 40.18 | 2 |

| | | | | | | |
|---|---|---|---|---|---|---|
| R12311 | 7.9 | 0 | M | AA | 58.48 | 2 |
| R12312 | 8.7 | 0 | M | AA | 48.66 | 2 |
| R12314 | 7.5 | 0 | F | AA | 59.58 | 2 |
| R12316 | 7.8 | 1 | F | AA | 48.93 | 2 |
| R12317 | 7.1 | 1 | F | CAUC | 71.81 | 2 |
| R12318 | 7.9 | 0 | M | AS | 58.2 | 2 |
| R12319 | 8 | 0 | F | AA | 67.73 | 2 |
| R12320 | 9 | 1 | M | AA | 46.47 | 2 |
| R12321 | 5.3 | 0 | F | AA | 44.14 | 2 |
| R12322 | 8.2 | 0 | M | AA | 51.59 | 2 |
| R12323 | 6.9 | 0 | F | CAUC | 51.91 | 2 |
| R12324 | 7.8 | 0 | F | CAUC | 64.28 | 2 |
| R12325 | 8.3 | 0 | F | AA | 36.48 | 2 |
| R12326 | 6.8 | 0 | M | AA | 60.35 | 2 |
| R12327 | 6.8 | 0 | M | AA | 68.54 | 2 |
| R12328 | 8.7 | 1 | F | CAUC | 44.86 | 2 |
| R12329 | 6.7 | 0 | M | AA | 32.98 | 2 |
| R12330 | 7.5 | 0 | M | AA | 18.48 | 2 |
| R12331 | 7.7 | 0 | F | AA | 53.86 | 2 |
| R12332 | 7.6 | 0 | M | AA | 60.7 | 2 |
| R12333 | 6.9 | 0 | M | AA | 73.23 | 2 |
| R12334 | 7.5 | 0 | F | AA | 48.1 | 2 |
| R12335 | 8.9 | 1 | F | CAUC | 26.07 | 2 |
| R12336 | 7.3 | 0 | F | AA | 54.48 | 2 |
| R12337 | 8.5 | 1 | M | CAUC | 24.33 | 2 |
| R12338 | 7.6 | 0 | M | CAUC | 61.13 | 2 |
| R12339 | 8 | 0 | F | AA | 41.17 | 2 |
| R12340 | 6 | 0 | M | AA | 49.72 | 2 |
| R12341 | 8.7 | 1 | F | AA | 48.26 | 2 |
| R12342 | 7.1 | 0 | M | CAUC | 49.18 | 2 |
| R12343 | 6.7 | 0 | F | AA | 51.8 | 2 |
| R12344 | 5.7 | 0 | F | CAUC | 57.44 | 2 |
| R12345 | 7.5 | 0 | M | CAUC | 46.53 | 2 |
| R12346 | 7.2 | 0 | M | AA | 60.72 | 2 |
| R12347 | 6.9 | 0 | M | CAUC | 18.41 | 2 |
| R12348 | 6.8 | 1 | F | AA | 62.39 | 2 |
| R12349 | 7.5 | 0 | M | AA | 58.19 | 2 |
| R12350 | 6.8 | 0 | M | CAUC | 47.45 | 2 |
| R12353 | 7.8 | 0 | M | AA | 62.74 | 2 |
| R12355 | 8.6 | 1 | F | AA | 42.98 | 2 |
| R12356 | 6.5 | 0 | M | AA | 58.4 | 2 |
| R12357 | 8.1 | 0 | F | AA | 53.36 | 2 |
| R12358 | 6.9 | 1 | F | AA | 63.05 | 2 |
| R12359 | 7.4 | 1 | F | AA | 57.13 | 2 |
| R12360 | 7.6 | 0 | F | AA | 38.59 | 2 |

| | | | | | | |
|---|---|---|---|---|---|---|
| R12361 | 6.8 | 0 | F | AA | 42.91 | 2 |
| R12362 | 7.5 | 0 | M | CAUC | 24.61 | 2 |
| R12363 | 7.8 | 0 | M | CAUC | 49.17 | 2 |
| R12364 | 8.3 | 1 | M | CAUC | 30.86 | 2 |
| R12365 | 7.9 | 1 | M | AA | 65.32 | 2 |
| R12366 | 7.2 | 0 | M | CAUC | 41.01 | 2 |
| R12367 | 8.4 | 1 | M | CAUC | 58.96 | 2 |
| R12368 | 7.5 | 0 | M | CAUC | 28.35 | 2 |
| R12369 | 8.2 | 1 | F | AA | 45 | 2 |
| R12372 | 7.1 | 0 | M | AA | 51.55 | 2 |
| R12373 | 8 | 1 | M | CAUC | 48.08 | 2 |
| R12377 | 6.7 | 1 | M | AA | 53.24 | 2 |
| R12378 | 6.8 | 1 | M | AA | 18.57 | 2 |
| R12379 | 7.6 | 1 | M | AA | 21.54 | 2 |
| R12380 | 8.1 | 0 | F | HISP | 56.09 | 2 |
| R12381 | 8.3 | 0 | M | CAUC | 34.23 | 2 |
| R12382 | 6.9 | 0 | F | AA | 40.82 | 2 |
| R12383 | 5.2 | 1 | F | AA | 55.69 | 2 |
| R12384 | 7 | 0 | F | CAUC | 38.39 | 2 |
| R12386 | 7.3 | 1 | M | AA | 48.34 | 2 |
| R12387 | 7.3 | 0 | F | AA | 65 | 2 |
| R12389 | 7.2 | 1 | F | CAUC | 92.1 | 2 |
| R12390 | 7.6 | 1 | F | CAUC | 96.92 | 2 |
| R12391 | 7.6 | 1 | F | AA | 71.12 | 2 |
| R12392 | 7.6 | 0 | M | AA | 64.52 | 2 |
| R12393 | 6.9 | 1 | F | AS | 57.69 | 2 |
| R12394 | 7.7 | 1 | M | CAUC | 30.97 | 2 |
| R12395 | 8.1 | 1 | M | AA | 38.6 | 2 |
| R12396 | 7.7 | 0 | F | CAUC | 30.02 | 2 |
| R12398 | 7.5 | 0 | F | AA | 64.86 | 2 |
| R12399 | 7.2 | 0 | F | CAUC | 19.26 | 2 |
| R12400 | 10 | 0 | F | AA | 46.07 | 2 |
| R12401 | 7.4 | 1 | F | AA | 62.68 | 2 |
| R12402 | 5.5 | 1 | F | AA | 52.13 | 2 |
| R12403 | 7.3 | 1 | M | CAUC | 70.24 | 2 |
| R12404 | 10 | 0 | M | CAUC | 46.57 | 2 |
| R12405 | 7.8 | 0 | M | AA | 53.26 | 2 |
| R12406 | 5.4 | 0 | F | CAUC | 61.15 | 2 |
| R12407 | 7.1 | 0 | F | AA | 54.56 | 2 |
| R12408 | 7.8 | 0 | M | CAUC | 20.85 | 2 |
| R12409 | 6.8 | 0 | M | CAUC | 51 | 2 |
| R12410 | 6.7 | 0 | M | CAUC | 39.64 | 2 |
| R12411 | 6.9 | 1 | M | AA | 57.58 | 2 |
| R12412 | 7.4 | 1 | M | AA | 37.67 | 2 |
| R12413 | 7.1 | 1 | F | CAUC | 56.91 | 2 |

| | | | | | | |
|---|---|---|---|---|---|---|
| R12414 | 7.9 | 0 | M | CAUC | 46.08 | 2 |
| R12415 | 5.3 | 0 | F | CAUC | 33.97 | 2 |
| R12416 | 5.3 | 1 | M | AA | 50.57 | 2 |
| R12417 | 6 | 1 | M | AA | 83.41 | 2 |
| R12418 | 6.5 | 1 | M | CAUC | 68.97 | 2 |
| R12419 | 5.7 | 1 | F | CAUC | 54.53 | 2 |
| R12420 | 6.3 | 1 | F | CAUC | 83.14 | 2 |
| R12421 | 7.8 | 0 | M | AA | 27.2 | 2 |
| R12422 | 7.5 | 1 | M | AA | 47.59 | 2 |
| R12423 | 7.8 | 0 | F | AA | 53.56 | 2 |
| R12424 | 7.6 | 0 | M | CAUC | 77.94 | 2 |
| R12425 | 7.5 | 0 | F | AA | 71.77 | 2 |
| R12426 | 7.9 | 1 | M | CAUC | 46.19 | 2 |
| R12427 | 7.6 | 0 | M | CAUC | 46.94 | 2 |
| R12428 | 7.6 | 0 | M | CAUC | 62.27 | 2 |
| R12429 | 8.3 | 1 | M | AA | 43.05 | 2 |
| R12430 | 8.1 | 1 | M | AA | 45.48 | 2 |
| R12431 | 7.7 | 1 | F | AA | 34.84 | 2 |
| R12432 | 8.4 | 1 | M | HISP | 48.81 | 2 |
| R12433 | 7.7 | 0 | F | CAUC | 83.59 | 2 |
| R12435 | 7.8 | 0 | M | CAUC | 42.81 | 2 |
| R12436 | 7.8 | 1 | M | CAUC | 68 | 2 |
| R12437 | 7.1 | 1 | M | CAUC | 42.71 | 2 |
| R12438 | 8 | 1 | M | CAUC | 41.33 | 2 |
| R12439 | 8.5 | 1 | M | AA | 52.02 | 2 |
| R12440 | 7.4 | 1 | M | HISP | 79.32 | 2 |
| R12441 | 7.5 | 1 | M | AA | 70.91 | 2 |
| R12442 | 7.5 | 0 | M | CAUC | 63.2 | 2 |
| R12443 | 6.8 | 0 | M | CAUC | 34.32 | 2 |
| R12444 | 7.6 | 1 | M | AA | 48.51 | 2 |
| R12445 | 8 | 1 | M | AS | 45.59 | 2 |
| R12446 | 5.3 | 1 | M | AA | 53.18 | 2 |
| R12447 | 7.6 | 0 | F | CAUC | 48.99 | 2 |
| R12448 | 7.8 | 0 | M | CAUC | 56.05 | 2 |
| R12449 | 7.2 | 0 | M | AA | 54.86 | 2 |
| R12450 | 7.8 | 1 | M | CAUC | 22.1 | 2 |
| R12456 | 8.3 | 0 | M | AA | 45.16 | 2 |
| R12457 | 7.3 | 0 | M | AA | 38.48 | 2 |
| R12458 | 8.6 | 0 | M | AA | 34.76 | 2 |
| R12460 | 8.4 | 0 | M | AA | 46 | 2 |
| R12461 | 7.8 | 0 | M | AA | 23.58 | 2 |
| R12462 | 7.4 | 0 | F | AA | 46.71 | 2 |
| R12463 | 7.1 | 0 | M | AA | 49.65 | 2 |
| R12464 | 6.9 | 0 | M | CAUC | 52.49 | 2 |
| R12465 | 8 | 1 | F | CAUC | 41.09 | 2 |

| | | | | | | |
|---|---|---|---|---|---|---|
| R12466 | 7.8 | 0 | M | CAUC | 55.51 | 2 |
| R12470 | 7.9 | 0 | M | AA | 29.06 | 2 |
| R12472 | 8.1 | 1 | M | CAUC | 62.2 | 2 |
| R12473 | 6.6 | 1 | M | AA | 59.06 | 2 |
| R12474 | 7.5 | 1 | M | AA | 50.34 | 2 |
| R12475 | 6.9 | 1 | F | CAUC | 57.04 | 2 |
| R12477 | 8 | 0 | M | AA | 73.86 | 2 |
| R12478 | 7.3 | 1 | M | CAUC | 55.68 | 2 |
| R12479 | 6.2 | 1 | M | CAUC | 56.15 | 2 |
| R12480 | 7.3 | 0 | M | CAUC | 55.91 | 2 |
| R12481 | 8.3 | 0 | M | CAUC | 40.08 | 2 |
| R12482 | 7.1 | 0 | M | AA | 65.58 | 2 |
| R12483 | 7.5 | 0 | M | AA | 40.24 | 2 |
| R12485 | 7.8 | 0 | M | AA | 56.48 | 2 |
| R12486 | 7.9 | 0 | M | CAUC | 36.53 | 2 |
| R12487 | 5.4 | 0 | M | CAUC | 48.07 | 2 |
| R12488 | 7.1 | 1 | M | AA | 49.26 | 2 |
| R12489 | 6 | 1 | M | AA | 30.68 | 2 |
| R12490 | 7.1 | 1 | F | CAUC | 50.73 | 2 |
| R12491 | 7.1 | 0 | M | CAUC | 59.22 | 2 |
| R12493 | 7.3 | 0 | F | AA | 40.31 | 2 |
| R12494 | 6.1 | 0 | M | AA | 49.39 | 2 |
| R12495 | 6.9 | 1 | M | CAUC | 50.8 | 2 |
| R12496 | 7.9 | 0 | M | CAUC | 53.29 | 2 |
| R12497 | 7.5 | 0 | F | AA | 50.84 | 2 |
| R12498 | 7.5 | 0 | M | CAUC | 44.7 | 2 |
| R12499 | 6.3 | 1 | M | CAUC | 20.34 | 2 |
| R12501 | 6.2 | 0 | M | AA | 57.82 | 2 |
| R12503 | 7.5 | 0 | M | CAUC | 63.5 | 2 |
| R12505 | 5.1 | 0 | F | CAUC | 62.99 | 2 |
| R12506 | 5.1 | 1 | F | CAUC | 59.93 | 2 |
| R12508 | 7.5 | 1 | M | CAUC | 54.73 | 2 |
| R12510 | 7.8 | 0 | M | CAUC | 23.22 | 2 |
| R12511 | 7.8 | 0 | F | AA | 56.6 | 2 |
| R12512 | 7.7 | 1 | M | CAUC | 38.41 | 2 |
| R12515 | 6.8 | 1 | F | AA | 66.01 | 2 |
| R12516 | 7.4 | 0 | F | AA | 35.8 | 2 |
| R12517 | 6.7 | 0 | M | CAUC | 56.82 | 2 |
| R12519 | 7.6 | 0 | M | HISP | 36.11 | 2 |
| R12520 | 7.5 | 0 | M | AS | 27.74 | 2 |
| R12521 | 6.8 | 0 | F | AA | 48.57 | 2 |
| R12522 | 6.7 | 0 | F | CAUC | 60.51 | 2 |
| R12523 | 6.7 | 1 | M | CAUC | 45.76 | 2 |
| R12524 | 6.1 | 1 | F | CAUC | 47.56 | 2 |
| R12525 | 7.6 | 0 | M | CAUC | 59.48 | 2 |

| | | | | | | |
|---|---|---|---|---|---|---|
| R12526 | 8 | 0 | M | CAUC | 46.12 | 2 |
| R12527 | 6.9 | 1 | M | AA | 53.95 | 2 |
| R12530 | 7.3 | 1 | M | AA | 53.74 | 2 |
| R12531 | 7.7 | 0 | M | CAUC | 51.05 | 2 |
| R12533 | 7.8 | 0 | M | CAUC | 52.33 | 2 |
| R12535 | 7.2 | 0 | M | AA | 84.16 | 2 |
| R12536 | 7.6 | 0 | M | CAUC | 54.9 | 2 |
| R12537 | 8.6 | 0 | M | AA | 50.47 | 2 |
| R12538 | 7.2 | 0 | M | CAUC | 40.38 | 2 |
| R12539 | 7.3 | 0 | M | CAUC | 41.44 | 2 |
| R12540 | 7.5 | 1 | F | AA | 33.35 | 2 |
| R12551 | 8.2 | 0 | M | CAUC | 63.16 | 2 |
| R12553 | 8.3 | 0 | M | CAUC | 42.05 | 2 |
| R12554 | 8.3 | 0 | M | CAUC | 51.23 | 2 |
| R12555 | 8.1 | 0 | M | CAUC | 51.63 | 2 |
| R12557 | 8.1 | 0 | F | CAUC | 57.83 | 2 |
| R12558 | 5.1 | 0 | F | AA | 50.6 | 2 |
| R12560 | 6.9 | 0 | M | CAUC | 47.83 | 2 |
| R12561 | 7.4 | 0 | M | CAUC | 72.43 | 2 |
| R12562 | 8.2 | 0 | M | AA | 71.4 | 2 |
| R12563 | 7 | 0 | M | CAUC | 61.75 | 2 |
| R12564 | 7.4 | 0 | M | HISP | 33.28 | 2 |
| R12565 | 8.1 | 0 | F | CAUC | 50.42 | 2 |
| R12566 | 7.4 | 0 | F | AA | 43.71 | 2 |
| R12567 | 8.2 | 0 | M | CAUC | 40.08 | 2 |
| R12568 | 8.3 | 0 | M | CAUC | 34.56 | 2 |
| R12569 | 8 | 0 | F | CAUC | 48.69 | 2 |
| R12570 | 8.1 | 0 | M | AA | 34.57 | 2 |
| R2816 | 8.4 | 1 | F | AA | 60.7 | 1 |
| R2824 | 6.2 | 0 | M | AS | 33.22 | 1 |
| R2867 | 8.5 | 1 | F | AA | 59.58 | 1 |
| R2905 | 8.6 | 0 | M | AA | 22.58 | 1 |
| R2947 | 8.7 | 0 | M | AA | 31.6 | 1 |
| R3002 | 8.1 | 0 | F | AA | 26.77 | 1 |
| R3029 | 8.3 | 0 | M | HISP | 48.47 | 1 |
| R3030 | 8.3 | 1 | F | AA | 66.29 | 1 |
| R3036 | 9 | 1 | M | AA | 75.57 | 1 |
| R3050 | 7.2 | 0 | M | AA | 21.01 | 1 |
| R3070 | 7.9 | 1 | M | CAUC | 81.16 | 1 |
| R3393 | 8.4 | 0 | M | AA | 52.13 | 1 |
| R3417 | 8.8 | 0 | M | AA | 19.89 | 1 |
| R3456 | 8.9 | 0 | M | AA | 24.25 | 1 |
| R3469 | 8.8 | 0 | M | AA | 45.86 | 1 |
| R3470 | 8.8 | 0 | M | AA | 18.28 | 1 |
| R3491 | 8.5 | 0 | M | CAUC | 32.67 | 1 |

| | | | | | | |
|---|---|---|---|---|---|---|
| R3496 | 8.9 | 1 | M | CAUC | 39.18 | 1 |
| R3517 | 8.5 | 0 | F | AA | 42.68 | 1 |
| R3540 | 8.2 | 1 | M | AA | 68.81 | 1 |
| R3555 | 9 | 1 | M | CAUC | 66.81 | 1 |
| R3557 | 9 | 0 | M | CAUC | 18.11 | 1 |
| R3575 | 9.1 | 1 | M | AA | 37.3 | 1 |
| R3597 | 8.7 | 0 | M | AA | 20.77 | 1 |
| R3606 | 9.3 | 1 | M | AA | 35.46 | 1 |
| R3611 | 8.8 | 1 | M | AA | 41.25 | 1 |
| R3614 | 7.4 | 1 | F | CAUC | 60.08 | 1 |
| R3616 | 8.8 | 1 | M | CAUC | 37.16 | 1 |
| R3639 | 8.7 | 0 | M | CAUC | 49.53 | 1 |
| R3645 | 5.9 | 1 | F | AA | 59.49 | 1 |
| R3646 | 7.2 | 1 | M | AA | 49.35 | 1 |
| R3665 | 8.4 | 0 | M | CAUC | 42.87 | 1 |
| R3670 | 8.3 | 1 | M | CAUC | 20.22 | 1 |
| R3774 | 8.8 | 0 | M | CAUC | 30.01 | 1 |
| R3887 | 8.7 | 0 | M | AA | 22.16 | 1 |
| R3888 | 6.9 | 1 | F | AA | 64.57 | 1 |
| R3891 | 8.8 | 0 | M | HISP | 19.38 | 1 |
| R3894 | 8.6 | 0 | F | AA | 31.53 | 1 |
| R3900 | 6.8 | 1 | F | CAUC | 59.82 | 1 |
| R3901 | 8.3 | 0 | M | CAUC | 50.87 | 1 |
| R3907 | 8.6 | 0 | F | AS | 78.18 | 1 |
| R3910 | 7.2 | 1 | F | CAUC | 32.01 | 1 |
| R3927 | 8.1 | 0 | F | AA | 45.38 | 1 |
| R3929 | 8.3 | 1 | M | AA | 60.11 | 1 |
| R3932 | 8.7 | 0 | M | CAUC | 53.94 | 1 |
| R3939 | 9.1 | 0 | F | AA | 29.95 | 1 |
| R3944 | 7.9 | 0 | M | AA | 45.29 | 1 |
| R3977 | 8.8 | 1 | M | AA | 60 | 1 |
| R3978 | 7.6 | 1 | M | CAUC | 55.68 | 1 |
| R4043 | 8.8 | 1 | M | AA | 51.71 | 1 |
| R4046 | 8.2 | 1 | M | CAUC | 55.85 | 1 |
| R4047 | 8.3 | 0 | M | CAUC | 33.24 | 1 |
| R4108 | 9.4 | 1 | M | AA | 53.12 | 1 |
| R4109 | 6.7 | 1 | M | AA | 74.56 | 1 |
| R4111 | 8 | 1 | M | AA | 43.35 | 1 |
| R4114 | 8.8 | 0 | M | CAUC | 28.57 | 1 |
| R4162 | 5.9 | 1 | M | AA | 61.46 | 1 |
| R4177 | 8.1 | 0 | M | AA | 22.71 | 1 |
| R4179 | 8 | 0 | M | AA | 35.24 | 1 |
| R4181 | 8 | 0 | F | AA | 21.53 | 1 |
| R4194 | 8.3 | 0 | M | HISP | 20.21 | 1 |
| R4200 | 7.8 | 0 | M | AA | 25.82 | 1 |

| | | | | | | |
|---|---|---|---|---|---|---|
| R4207 | 7 | 0 | F | AA | 44.82 | 1 |
| R4226 | 8.9 | 0 | M | AA | 53.92 | 1 |
| R4273 | 8.2 | 1 | F | CAUC | 28.22 | 1 |
| R4279 | 9 | 0 | M | CAUC | 38.76 | 1 |
| R4338 | 8.3 | 0 | M | AS | 70.9 | 1 |
| R4339 | 8.4 | 1 | M | CAUC | 57.14 | 1 |
| R4351 | 5.8 | 1 | M | CAUC | 52.13 | 1 |
| R4355 | 7.4 | 1 | M | CAUC | 29.92 | 1 |
| R4357 | 8.1 | 1 | M | CAUC | 45.94 | 1 |
| R4364 | 8.3 | 1 | M | CAUC | 29.23 | 1 |
| R4366 | 7 | 1 | M | CAUC | 49.4 | 1 |
| R4367 | 7.2 | 1 | F | CAUC | 61.52 | 1 |
| R4381 | 8.7 | 1 | M | CAUC | 34.06 | 1 |
| R4383 | 8.5 | 1 | F | CAUC | 53.56 | 1 |
| R4387 | 8.3 | 1 | M | CAUC | 28.59 | 1 |
| R4388 | 6.4 | 1 | F | AA | 45.73 | 1 |
| R4393 | 8.5 | 1 | M | CAUC | 21.24 | 1 |
| R4407 | 8.5 | 1 | F | CAUC | 53.39 | 1 |
| R4414 | 8.2 | 1 | M | CAUC | 18 | 1 |
| R4417 | 7.8 | 1 | F | AA | 52.22 | 1 |
| R5600 | 8.7 | 1 | M | AA | 44.64 | 1 |
| R5608 | 7.1 | 1 | F | CAUC | 46.53 | 1 |
| R5618 | 7.4 | 1 | F | AA | 66.94 | 1 |
| R5622 | 8.7 | 0 | M | AA | 44.72 | 1 |
| R5743 | 8.8 | 0 | M | Multi-Racial | 52.54 | 1 |
| R5747 | 8.3 | 0 | M | HISP | 28.06 | 1 |
| R5853 | 7.9 | 0 | M | HISP | 35.54 | 1 |
| R5859 | 8.4 | 0 | F | AA | 21.32 | 1 |
| R5861 | 8.6 | 0 | M | AA | 48.61 | 1 |
| R5863 | 8 | 0 | M | AA | 49.73 | 1 |
| R5868 | 7.7 | 0 | M | AA | 48.59 | 1 |
| R5871 | 8.6 | 0 | M | CAUC | 18.72 | 1 |
| R5885 | 8.2 | 0 | M | CAUC | 51.72 | 1 |
| R5905 | 9.1 | 1 | M | CAUC | 53.97 | 1 |
| R6531 | 9.1 | 1 | M | AA | 52.89 | 1 |
| R6532 | 6.6 | 1 | M | CAUC | 19.69 | 1 |
| R6534 | 6.7 | 1 | M | CAUC | 21.06 | 1 |
| R6572 | 8.8 | 1 | M | AA | 70.52 | 1 |
| R6578 | 8.4 | 1 | M | CAUC | 57.58 | 1 |
| R6579 | 8.9 | 1 | M | CAUC | 60.56 | 1 |
